# Analysis of codon usage and allele frequencies reveal the double-edged nature of cross-kingdom RNAi

**DOI:** 10.1101/2022.07.19.500629

**Authors:** Bernhard T. Werner, Annette Kopp-Schneider, Karl-Heinz Kogel

## Abstract

**Background:** In recent years, a new class of small 21- to 24-nt-(s)RNAs has been discovered from microbial pathogens that interfere with their host’s gene expression during infection, reducing the host’s defence in a process called cross-kingdom RNA interference (ckRNAi). According to this model, microbial sRNAs should exert selection pressure on plants so that gene sequences that reduce complementarity to sRNAs are preferred. In this paper, we test this consequence of the ckRNA model by analyzing changes to target sequences considering codon usage and allele frequencies in the model system *Arabidopsis thaliana* (At) – *Hyaloperonospora arabidopsidis* (Ha) and *Hordeum vulgare* (Hv) – *Fusarium graminearum* (Fg). In both pathosystems, some selected sRNA and their corresponding target have been described and experimentally validated, while the lengthy methodology prevents the analysis of all discovered sRNAs. To expand the understanding of ckRNAi, we apply a new in silico approach that integrates the majority of sRNAs.

**Results:** We calculated the probability (P_CHS_) that synonymous host plant codons in a predicted sRNA target region would show the same or stronger complementarity as actually observed and compared this probability to sets of virtual analogous sRNAs. For the sets of Ha and Fg sRNAs, there was a significant difference in codon usage in their plant gene target regions (for Ha: P_CHS_ 24.9% lower than in the virtual sets; for Fg: P_CHS_ 19.3% lower than in the virtual sets), but unexpectedly for both sets of microbial sRNA we found a tendency towards codons with an unexpectedly high complementarity. To distinguish between complementarity caused by balancing sRNA-gene coevolution and directional selection we estimated Wright’s F-statistic (*F_ST_*), a measurement of population structure, in which positive deviations from the background indicate directional and negative deviations balancing selection at the respective loci. We found a negative correlation between P_CHS_ and *F_ST_* (p=0.03) in the At-Ha system indicating deviations from codon usage favoring complementarity are generally directionally selected.

**Conclusion:** The directional selection of complementary codons in host plants suggests an evolutionary pressure to facilitate silencing by exogenous microbial sRNAs, which is not consistent with the anticipated biological role of pathogen sRNAs as exclusively effectors in cross-kingdom RNAi. To resolve this conflict, we propose an extended model in which microbial sRNAs are perceived by plants via RNA interference and, via coevolution, primarily help to fine-tune plant gene expression.

## Background

Cross-kingdom RNA interference (ckRNAi) is a process in which small RNAs (sRNAs) are transferred from one organism to one from a different kingdom of life, where they cause gene silencing of complementary genes in the receiving organism (Baulcombe 2013). In the field of plant – microbe interactions, ckRNAi first came into focus through work by Hailing Jin’s group, Riverside. In a game changing work, they showed that the ascomycete fungal pathogen *Botrytis cinerea* (Bc) delivers 21 nucleotide (nt) sRNAs into its host plants *Arabidopsis thaliana* (At) and tomato (Weiberg et al. 2013; Wang et al. 2017a). Fungal sRNAs were shown to operate as RNA effector molecules that interfere with and silence plant defense genes such as mitogen-activated protein kinases *MPK1* and *MPK2*. This added another mosaic to our understanding of plant - pathogen interactions, as in the past effectors have typically been defined as mostly smaller, microbe-derived proteins that interfere with components of the plant immune system (He et al. 2020). According to the new understanding, sRNAs, like protein effectors, are the product of an evolutionary arms race between host and microbe (Jones and Dangl 2006).

In 2016, Hui-Shan Guo’s group also demonstrated the transfer of plant sRNAs into an interacting fungus: cotton plants export micro (mi)RNAs into the pathogenic ascomycete fungus *Verticillium dahliae*, and some of these miRNAs target fungal virulence genes (Zhang et al. 2016). Further work has shown that this exchange takes place via extracellular vesicles, which, in addition to other cellular components, also carry sRNAs as cargo (Cai et al. 2018a; Rutter and Innes 2018; Zand Karima et al. 2022; for review: Cai et al. 2021; Nasfi and Kogel 2022). Proof in principle of the transfer of sRNA from the host plant to a fungus had already been provided in 2010 by the discovery of the process of host-induced gene silencing (HIGS; Nowara et al. 2010). Transgenic barley plants producing double-stranded (ds)RNA exported corresponding sRNAs into the powdery mildew fungus, where they subsequently silenced target genes. Many reports have confirmed that HIGS can mediate strong resistance against target organisms including fungi, oomycetes and insects thereby demonstrating the great agronomic potential of artificial sRNAs (Knip et al. 2014; Cai et al. 2018b; Liu et al. 2020; Šečić and Kogel 2021).

In recent years many more sRNA effectors have been discovered in plant pathogenic fungi (Wang et al. 2017b; Dubey et al. 2019; Werner et al. 2021), oomycetes (Dunker et al. 2020), beneficial fungal endophytes (Šečić et al. 2021), and bacteria (Ren et al. 2019). Their function as effectors is often experimentally demonstrated by knock-out (KO) of *Dicer-like* (*DCLs*) genes in the microbe and/or *Argonaute* (*AGO*) genes in the host, as loss-of-function mutations correspondingly impair either the microbe’s ability to produce sRNAs or the host’s ability to recognize silencing signals (Weiberg et al. 2013; Cai et al. 2021), while arguably DCLs and AGOs also play important roles in virulence and immunity both in plants (Fang et al. 2016) and fungi (Nicolás et al. 2013, Gaffar et al. 2019), independent of ckRNAi.

Further experimental evidence for sRNA effector activity is provided by detecting corresponding degraded target host mRNAs fragments using degradome sequencing, also referred to as parallel analysis of RNA ends (PARE, German et al. 2008), and/or through recording their effect on pathogenicity upon artificial overexpression in the host (Weiberg et al. 2013). An elegant strategy to detect ckRNAi is the introduction of short tandem target mimics (STTM) that provides artificial RNA target sequences such that a cognate sRNA rather forms a complex with the target mimic, thus preventing silencing activity on its natural target (Yan et al. 2012; Zhang et al. 2016; Dunker et al. 2020). Dunker et al. (2020) examined plant *At*AGO1-linked sRNAs of the oomycete pathogen *Hyaloperonospora arabidopsidis* (Ha) after infection of Arabidopsis. Three of these sRNAs were confirmed as enhancing the pathogenicity of the oomycete by employing the STTM technique (Dunker et al. 2020). Above all, these recent discoveries have one key aspect in common: Plants and microbes can take up exogenous sRNA duplexes. This is at the first glance puzzling because as shown by the above examples uptake can have detrimental effects on the fitness and survival of the receiving organism.

Given the current knowledge of HIGS and ckRNAi, we speculated that plants might acquire synonymous mutations during the evolutionary arms race with microbial pathogens to evade silencing of key immune factors with relatively low fitness costs. Such an evolutionary strategy was shown to be feasible by Dunker et al. (2020), who demonstrated that sRNA-resistant versions of plant targeted genes, when generated by artificially introduced synonymous mutations, rendered host plants more resistant to Ha. Synonymous mutations occur due to the degenerative nature of the genetic code which enables organisms to encode the 20 possible amino acids (aa) with 64 three base long codons in mRNA (Crick et al. 1961). Even if these codons code for the same aa, there is an evolutionary cost attached to it and therefore these synonymous mutations are often non-silent. For example, At and other organisms prefer certain codons over others, especially in highly expressed genes, due in part to differences in translational efficiency, ultimately leading to a codon bias (Duret & Mouchiroud 1999).

An established approach for the detection of genes under selection is the analysis of allele distribution in distinct populations via Wright’s F-statistic (*F_ST_*) (Wright 1931). To this end, the distribution of *F_ST_* for distinct loci is compared to the background distribution of *F_ST_* in the genome (Akey et al. 2002). Positive outliers of *F_ST_* from the background are correlated to directional selection while negative outliers are correlated to sites under balancing selection (Akey et al. 2002, Beaumont & Balding 2004). For this kind of analysis, the 1001 genomes project, started in 2008, provides a solid foundation of data with 10.7 million single nucleotide polymorphisms (SNPs) and 1.4 million indels across 10 populations and 1135 individual At genomes (Alonso-Blanco et al. 2016).

Avoiding the formation of sRNA-mRNA complexes and gene silencing of defence genes is an advantage for the plant in ckRNAi systems. Here we tested the hypothesis that codon usage in target regions of sRNA effectors should differ from other coding regions in a plant genome, due to the selection of codons with lower complementarities to the respective pathogen sRNA. Additionally, because sRNA effectors potentially have a high plasticity, we expected a coevolution of effector and plant genes, where sRNAs complementary to the changed codon usage are selected for (Fig. 1). This process would lead to a reduced *F_ST_*, due to balancing selection, and exceptionally non-complementary codon usage.

**Figure 1:**
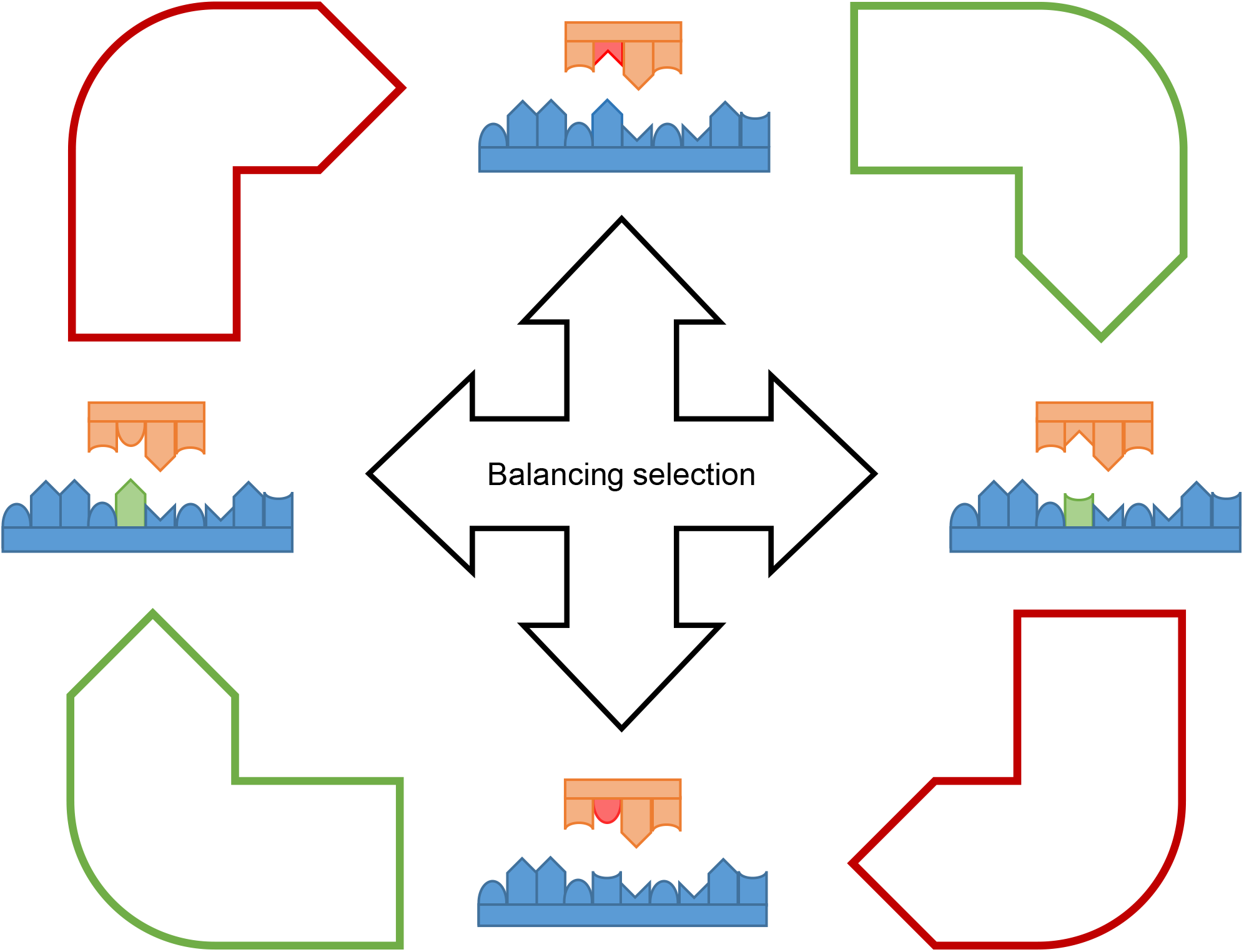
Mechanism of balancing selection between ckRNAi exerting sRNA and target mRNA under the effector hypothesis. mRNAs (blue) and sRNAs (orange) are shown stylized and shortened. Nucleic acids are depicted as shapes representing complementarity. Sequence changes are indicated with green (disruption of silencing) and red (restoration of silencing). Under the ckRNAi effector hypothesis silencing is expected to be detrimental for the sRNA receiving organism (e.g. the host plant) and beneficial for the sRNA sending organism (e.g. the pathogen). As a results of coevolution we expect all 4 states depicted to be present in the host and pathogen populations.

## Results

To reveal the difference in codon usage between sRNA target and non-target regions, we compared the observed bias in codon usage of sRNA complementary regions, which is predicted by the target prediction algorithm TAPIR (Bonnet et al. 2010), with the bias created by natural selection in response to ckRNAi. To achieve this, we compared the codon usage of published sets of At-miRNAs or microbial sRNAs from Ha and the plant pathogenic ascomycete *Fusarium graminearum* (Fg), respectively, to virtual sets of sRNAs (random (r)sRNAs). These rsRNAs were generated randomly by using the relative abundance of nucleotides (nts) in the respective published RNA set, to generate analogous sRNAs with the same length, size (Tab. S1 & S2) and base composition (Tab. S3) as the respective set of miRNAs or microbial sRNAs. Figure S1 gives an overview of our workflow.

For comparing the codon usage in the target regions of sRNAs and rsRNAs we calculated the probability P_CHS_ of each sRNA’s and rsRNA’s interaction with a coding sequence (CDS) having the same or a higher complementarity, under the assumptions that i. the aa sequence is conserved and ii. synonymous codons are chosen randomly, following the overall codon usage of the CDS of the host organism (Tab. S4 & S5). The procedure is pictured for At-miRNA400 as an example (Fig. 2). At-miRNA400 targets several pentatricopeptide repeat (PPR) proteins and is involved in biotic stress responses (Park et al. 2014). In Figure 1A the alignment of At-miRNA400 and two *PPR* genes is shown. The miRNA is perfectly complementary to *PPR1* (AT1G06580), while there are four mismatches (MM) to its homolog *T8K14.4* (AT1G79540). Remarkably, the underlying aa sequence of both PPRs is identical in the miRNA target site VTYNTLI.

**Figure 2:**
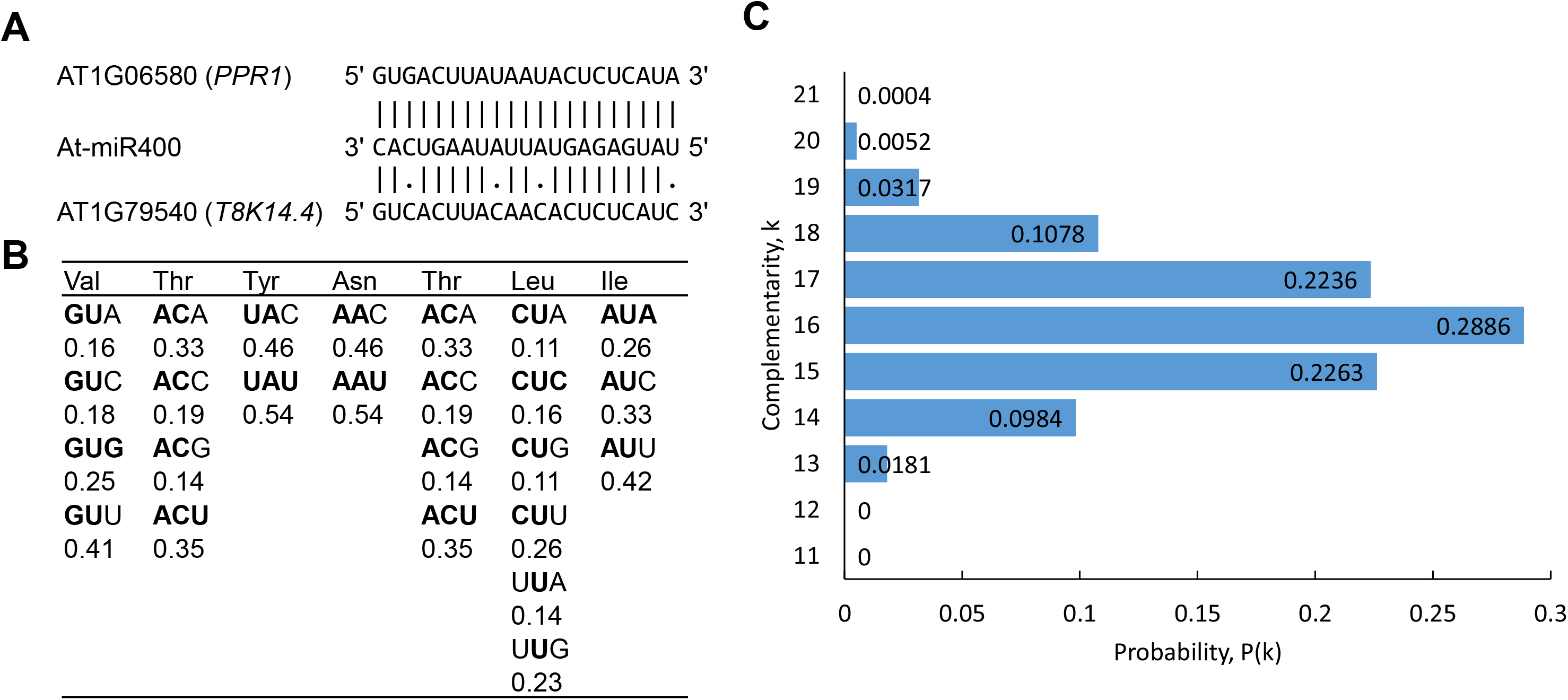
Exemplary calculation of the probability of a given complementarity between At-miRNA400 and the aa motif VTYNTLI present in PPR1 (AT1G06580) and T8K14.4 (AT1G79540). A: The alignment of AT1G06580 and AT1G79540 to At-miRNA400 is shown, with MM annotated as “.” and complementary bases as “|”. B: The distribution of synonymous codons for the aa present in the aa motif VTYNTLI in the At-CDS. Bases complementary to the miRNA in the respective position are marked in bold. C: By calculating the probability of all combinations of codons and the respective number of complementary bases the total probability for each number of complementary bases is shown.

The relative abundance for all synonymous codons for this aa motif is shown in Figure 1B, with the nts complementary to At-miRNA400 marked in bold. This data enables the calculation of the probability of different number of target site nts being complementary to the miRNA under assumptions i & ii (Fig. 1C) using the distribution of synonymous codons given in Figure 1B. For instance, the respective value for the 21 nts target site being fully complementary to At-miRNA400 is p = 0.25⋅0.35⋅0.54⋅0.54⋅0.35⋅0.16⋅0.26 = 0.0004, and the respective value for 17 nts is p = 0.2236, summing all probabilities for combining 4 MM in the 21 positions. To predict an interaction between sRNA and mRNA leading to silencing the TAPIR algorithm (Bonnet et al. 2010) applies a target score cutoff. This score is calculated by increasing the score for each mismatch by 1 and for each G-U alignment by 0.5 with doubled values in the seed region (nt 2-12). The four MM with one MM in the seed region between At-miRNA400 and *T8K14.4* lead to a target score of 5 which is above the typical threshold (target score ≤ 4) for silencing to occur in At (Bonnet et al. 2010). The probability (P_CHS_) for the same or a higher complementarity between miRNA400 is 0.0004 for *PPR1* and 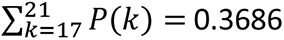 for *T8K14.4*. These P_CHS_ are consistent with the view that some PPR proteins being silenced during the expression of At-miRNA400 are beneficial for At, while the silencing of other PPR’s would be detrimental, and that for the beneficial case synonymous mutations were positively selected to enable this selective silencing of proteins carrying the same motif.

While this selection can be obvious in the case of PPR proteins, other genes with less family members should still be subject to the same kind of selective pressure where either silencing is beneficial or detrimental in the absence of another protein with an identical aa motif present in the genome. This pressure would persist until mutations lead to the abolishment of silencing. To establish a baseline probability, we predicted via TAPIR (Bonnet et al. 2010) all possible interactions and interactions one seed MM above the default threshold (target score ≤ 6) (Tab. S6) of all known At-miRNAs from miRBase (Griffiths-Jones et al. 2006) and a set of rsRNAs with the same size and nt abundances as the published At*-*miRNAs and calculated the P_CHS_ for all interactions in R (R Core Team 2021). We repeated the simulation experiment three times for three different sets of rsRNAs (Fig. 3A). The P_CHS_ of the At-miRNAs is significantly higher than the P_CHS_ of the rsRNA sets (estimated ratio of P_CHS_ and [95%-CI] : 1.204 [1.12, 1.295]; 1.112 [1.035, 1.194]; 1.226 [1.14, 1.319]). This could be the consequence of an adaptation of miRNAs to target-specific motifs consisting of the most common codons. Interestingly however, some interactions of the At-miRNAs have a very low P_CHS_, suggesting a change in codon usage to enable silencing as shown for the At-miRNA400 - *PPR1* interaction. To further substantiate our calculation, we repeated this experiment three times for the total set of *Gorilla gorilla* (Gg) miRNAs as a negative control (Fig. 3B). Consistent with the obvious lack of biological significance for Gg-miRNAs in the At-genome we did not see significant differences in the P_CHS_ for any set (estimated ratio of P_CHS_ and [95%-CI] : 1.034 [0.948, 1.127]; 1.058 [0.974, 1.15]; 1.039 [0.954, 1.131]).

**Figure 3:**
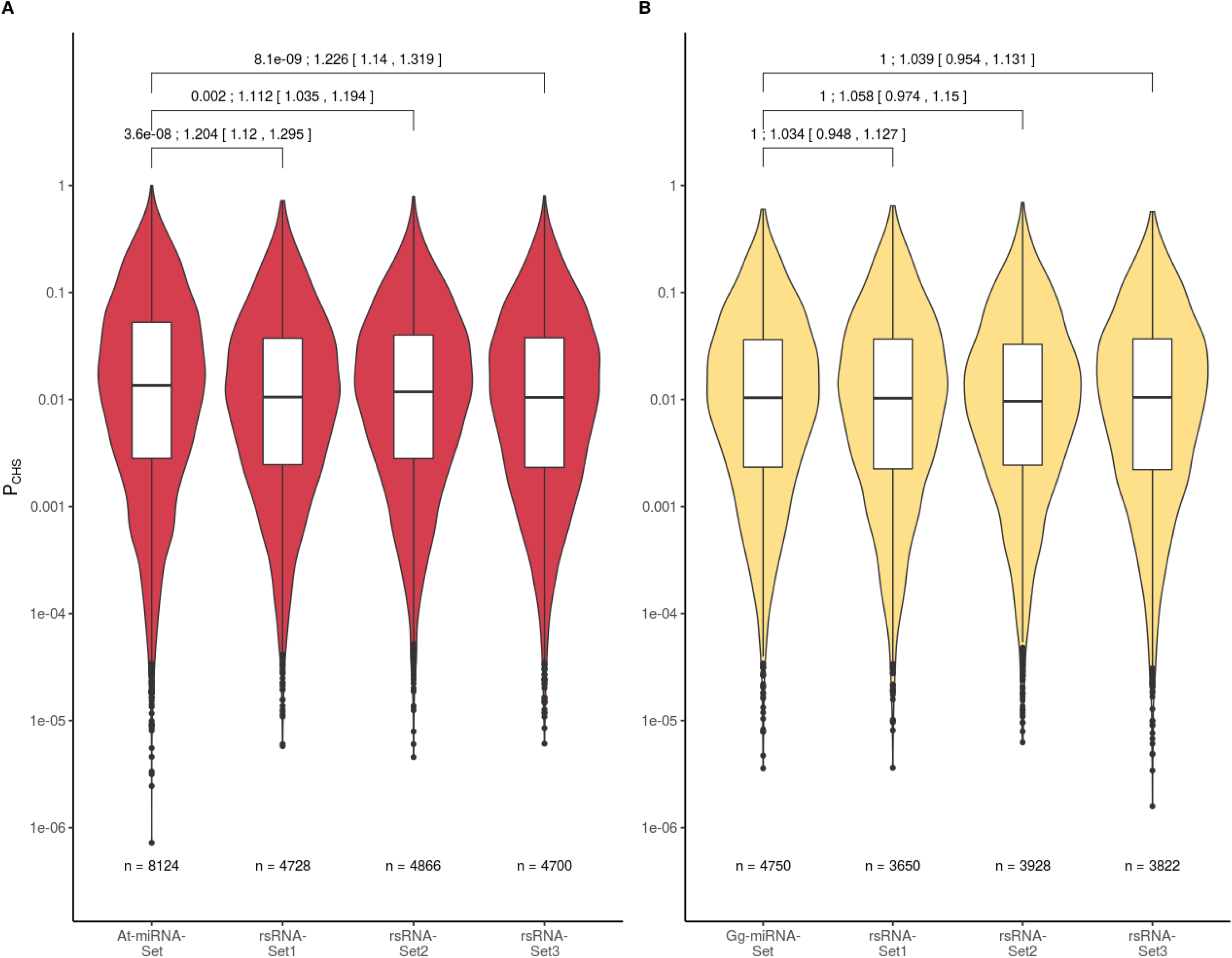
Calculated P_CHS_ values for the predicted mRNA-sRNA interactions in the At-CDS by At- and Gg-miRNAs. The violin plots with internal boxplots show the distribution of P_CHS_ values of the computationally predicted mRNA-sRNA interactions. The P_CHS_ is the probability of a random selection of synonymous codons based on the relative frequency of codons in the At-CDS to have the same complementarity or a higher complementarity as the actual complementarity. The values are shown for different sRNA sets taken from miRBase, A: At-miRNAs; B: Gg-miRNAs as a negative control, and three analogous random sets of sRNAs (rsRNA-sets, see methods). p-values were calculated with a Kruskal-Wallis test adjusted for multiple testing after Benjamini & Yekutieli (2001). Only p values for comparisons between published and rsRNA sets are shown. There were no significant differences between individual rsRNA-sets. To achieve normality, P_CHS_ values were log_10_-transformed. The 95% confidence interval (95%-CI) for the difference of mean log_10_(P_CHS_) was calculated via a two-sample Welch t-test and subsequently retransformed and is shown as relative difference to the respective rsRNA-set. n refers to the total number of predicted interactions. The results of the statistical tests are shown as: p-value; arithmetic mean [upper and lower bounds of 95%CI]

In a next step, we applied the same methodology established for At-miRNA to the microbial Ha-sRNA set provided by Dunker et al. (2020). We analyzed all sRNAs with at least an average of 100 reads per million in the published datasets (Fig. 4A & Tab. S1). In all three repetitions, the median P_CHS_ was lower for the Ha-sRNAs compared to the respective rsRNA-sets, showing that target sites in the At-genome for Ha-sRNAs, like At-miRNAs, evolved a significantly different codon usage, supporting our hypothesis. Unexpectedly, however, the codon usage in Ha-sRNAs target regions was biased towards codons with an surprisingly high complementarity to the pathogen sRNAs, reflected in a lower P_CHS_ (estimated ratio of P_CHS_ and [95%-CI]: 0.62 [0.472, 0.814]; 0.758 [0.576, 0.998]; 0.595 [0.447, 0.793]). Next, we analyzed a published sRNA dataset from *Fusarium graminearum* (Werner et al. 2021) in combination with the CDS from its host plant barley (*Hordeum vulgare*). Fg is an important plant pathogen with identified sRNA-effectors (Werner et al. 2021; Jian & Liang 2019). Again, we found that the Fg-sRNAs targeted regions were composed of codons with an unlikely high complementarity to the respective sRNA, compared to the rsRNAs (estimated ratio of P_CHS_ and [95%-CI]: 0.711 [0.639, 0.791]; 0.732 [0.66, 0.813]; 0.801 [0.721, 0.89]) (Fig. 4B). This bias in host plants suggests an evolutionary pressure to facilitate silencing by exogenous RNAs, which challenges the conjectured role of pathogen sRNAs as primarily effector-like. To resolve this contradiction, we propose an extended model, in which microbial sRNAs are perceived by plants via RNAi, and predominately aid in the fine-tuning of plant gene expression, while the role of sRNAs as effector is less prevalent.

**Figure 4:**
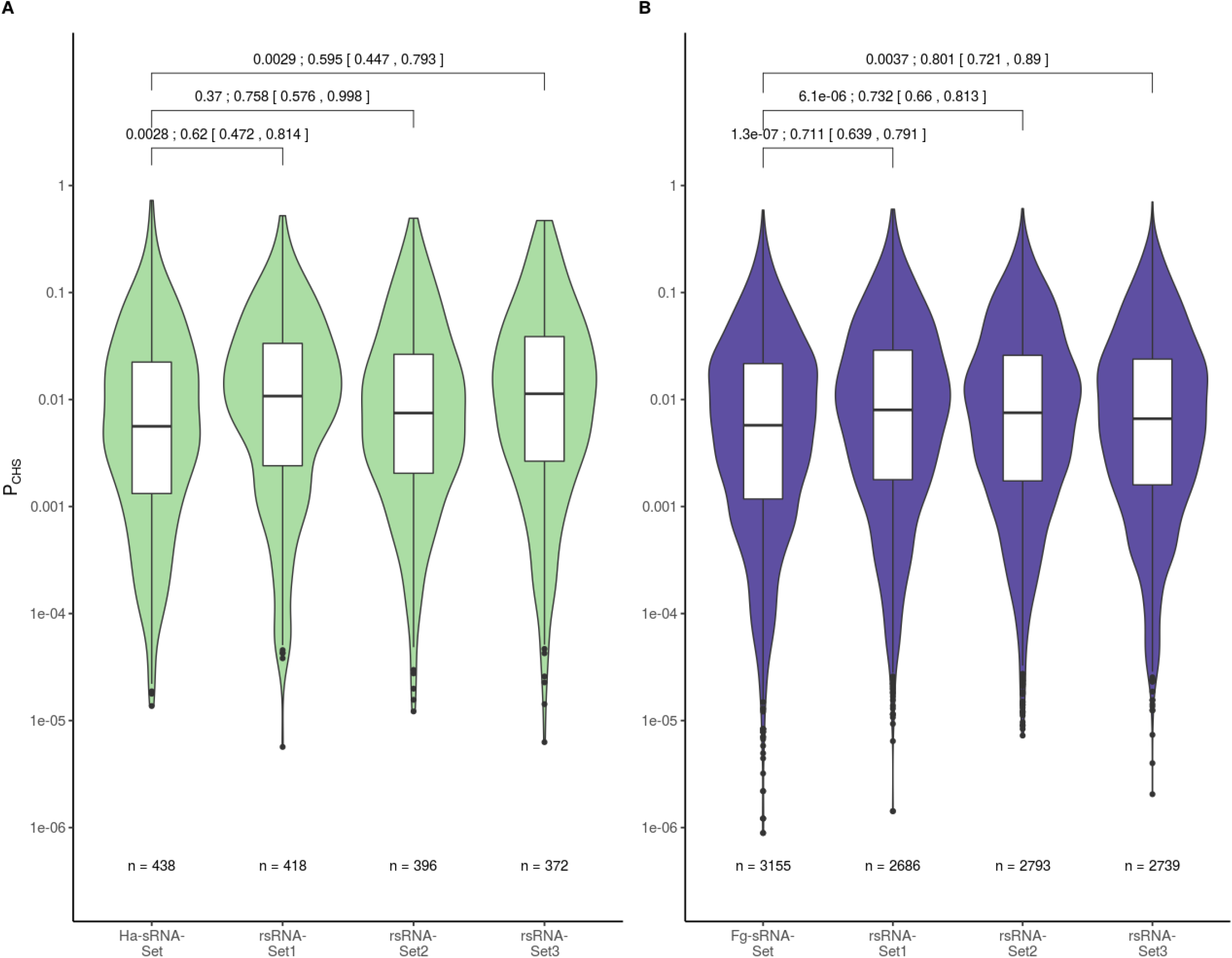
Calculated P_CHS_ values for the predicted mRNA-sRNA interactions in the At-CDS with Ha*-* sRNAs and barley-CDS with Fg*-*sRNAs. The violin plots with internal boxplots show the distribution of P_CHS_ values of the computationally predicted mRNA-sRNA interactions. The P_CHS_ is the probability of a random selection of synonymous codons based on the relative frequency of codons in the At- or Hv-CDS to have the same complementarity or a higher complementarity as the actual complementarity. The values are shown for sRNA-sets from two organisms A: Ha-sRNAs from an At-AGO1 co-IP experiment (Dunker et al. 2020) and B: Fg-sRNAs from axenic culture (Werner et al. 2021) and three virtual analogous random sets of sRNAs (rsRNA-sets). p-values were calculated with a Kruskal-Wallis test adjusted for multiple testing after Benjamini & Yekutieli (2001). Only p values for comparisons between published and rsRNA sets are shown. There were no significant differences between individual rsRNA-sets. To achieve normality, P_CHS_ values were log_10_-transformed. The 95% confidence interval (95%-CI) for the difference of mean log_10_(P_CHS_) was calculated via a two-sample Welch t-test and subsequently retransformed and is shown as relative difference to the respective rsRNA-set. n refers to the total number of predicted interactions. The results of the statistical tests are shown as: p-value; arithmetic mean [upper and lower bounds of 95%CI]

Since 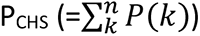 is increasing monotonously with decreasing complementarity k, i.e. with increasing number of mismatches between sRNA and mRNA, we wanted to exclude the possibility that an evolution of pathogen sRNAs leads to a lower number of mismatches between sRNA and target causing the observed lower P_CHS_ for interactions of pathogen sRNAs in contrast to rsRNAs. Therefore, we analyzed the target regions of our sRNA-sets for signs of balancing selection, because in such a scenario of sRNA-mRNA coevolution we would expect this kind of selection. To this end we utilized the data generated by the 1001 genomes project (Alonso-Blanco et al. 2016). This dataset with its high density of high confidence SNPs (10.7 million SNPs in 119 Mb genome) and relatively high sample sizes (1135 genomes) enables the analysis of the relatively short target sequences of individual sRNAs (21-24 base pairs) of Ha in At.

The 1135 genomes are assigned to 9 populations with admixed individuals assigned by Alonso-Blanco et al. (2016). We depicted the geographical distribution and the respective population of individuals in Figure 5 with the priorly analyzed line Col-0 marked, which belongs to the German population. Col-0 is the major isolate for At research being the first fully sequenced plant with a vast library of KO-mutants available. Due to the necessity of a high humidity for the life cycle of Ha we analyzed weather data from the closest weather stations to the location of individual At isolates to select a suitable subpopulation for the calculation of single locus *F_ST_*. To this end we analyzed data from the Integrated Surface Database (ISD) (Smith et al. 2011) provided by the National Oceanic and Atmospheric Administration (NOAA) with the R package rnoaa (Chamberlain 2021). Stations were selected for continuous operation from 1986 – 2016, to estimate the climate for each population, while we restricted the collected data to every third year to reduce the amount of hourly observations to a manageable amount. Figure 6 shows each weather station in our analysis in the European region for readability, while the map for all stations is shown in Figure S2.

**Figure 5:**
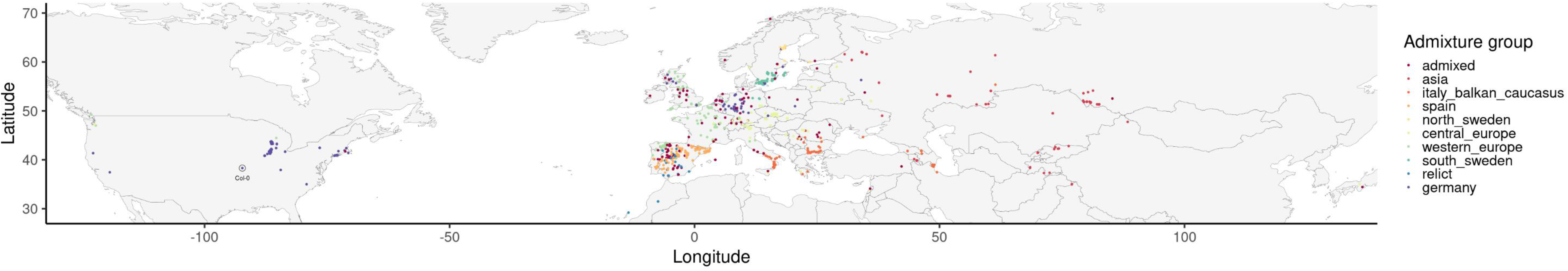
Sampling location and population origin of all At accessions from the 1001 genome project. This map in equirectangular projection shows the sample location of all lines present in the Arabidopsis 1001 genomes project. Each dot represents one line with colors indicating the respective population. Admixed lines are shown in red. The populations colors are ordered to enable maximum distinction between populations.

**Figure 6:**
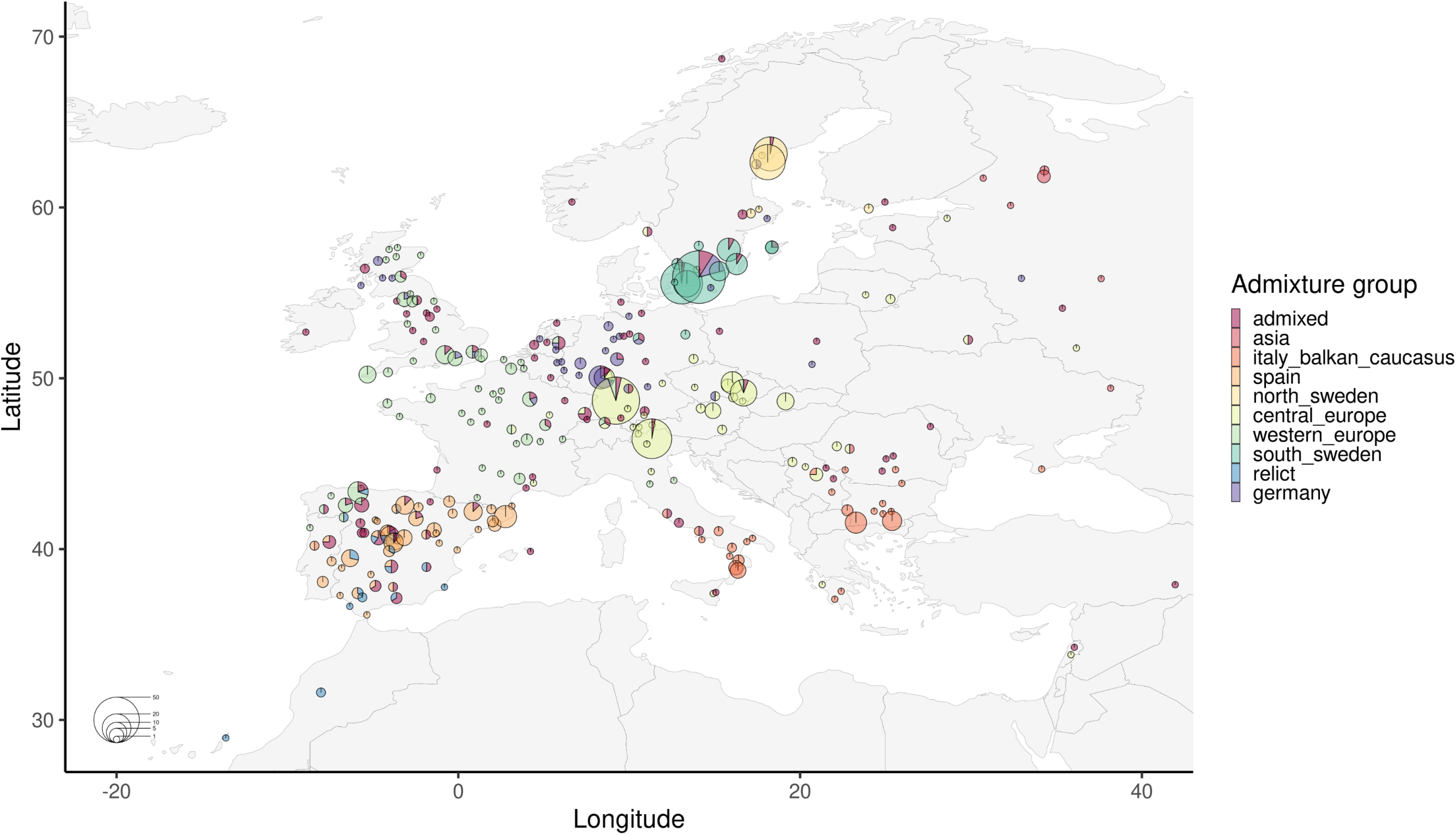
Weather stations used for the climate estimates of At populations. This equirectangular map of Europe shows locations of all European weather stations used to estimate the climatic conditions for each population. Each pie chart represents one station. Size indicates the number of lines represented by the station and colors indicate the respective population of lines represented.

Optimal conditions for downy mildew pathogenesis are widely researched and differ with developmental stages and species. Depending on developmental stage optimal temperatures are reported as generally 10-15°C, for germination of conidia 8-12°C, for penetration 16°C and for haustoria formation 20-24°C for Ha (Slusarenko & Schlaich 2003). Other authors report optimal temperatures for disease development at 20°C with no development of infection above 35°C (Achar 1998). For other species of the genus Peronospora infectivity is reported between 0-30°C with bimodal optima in the range of 10-25°C (Arauz et al. 2010; Choudhury & McRoberts 2017). While temperature constraints are less severe for the development of downy mildew infections high humidity is essential. The priory mentioned research on temperature constraints were conducted at 100% relative humidity (RH). A study series conducted by Hartmann et al. (1982; 1983) established a limit for Ha germination between -30 --60 bar atmospheric water potential which is equivalent to a RH ∼>94%. Therefore, we calculated the relative number of days with at least one measurement at the closest weather station with tempratures between 0-30°C and a RH >94% for each isolate in every third year between 1986 – 2016.

The highest contrast in days with potential Ha infection is between the German and south Swedish populations with 22% vs. 59% (Fig. 7). The low Ha potential for the American isolates (Fig. S3) contributes to this strong contrast, yet the similar sample sizes of 171 vs. 152 lines and the clear difference in Ha potential between the historical habitat of the two populations makes the south Swedish population the ideal candidate for the analysis of allele frequencies.

**Figure 7:**
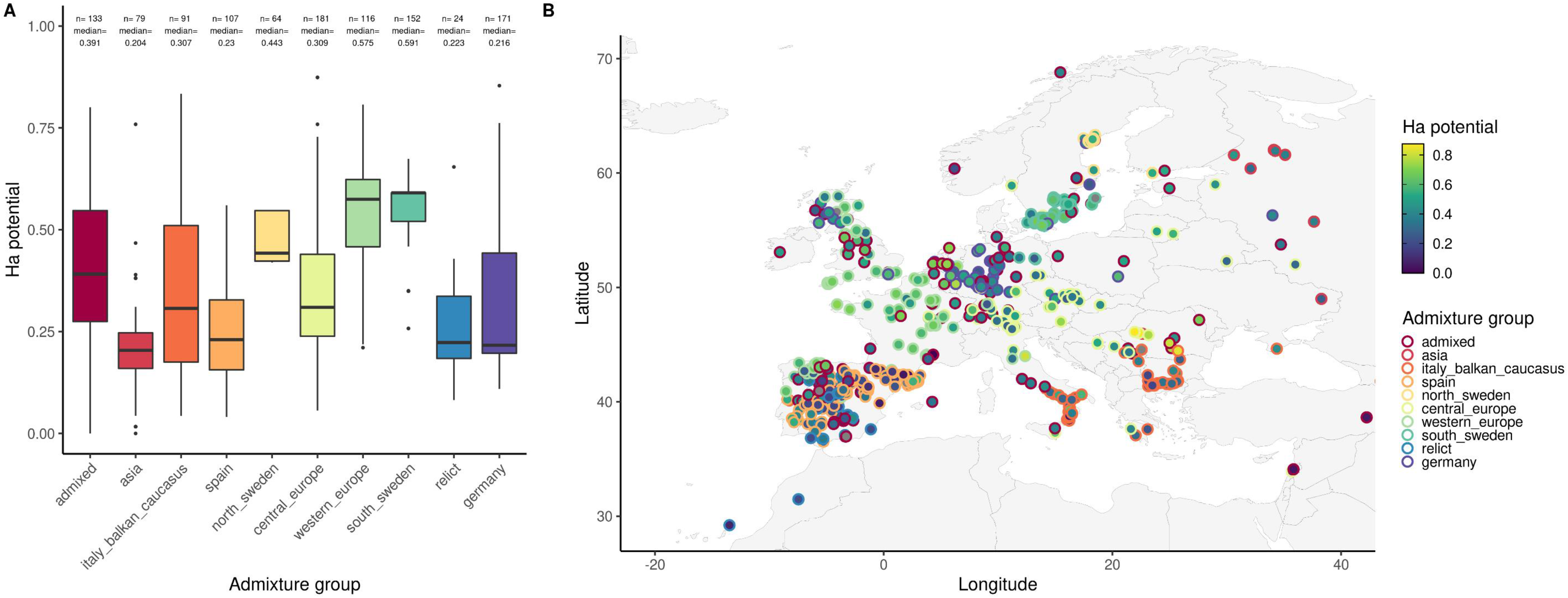
Estimated potential for Ha infections based on climate data. The potential for Ha infections was estimated by the analysis of weather data from the Integrated Surface Database (ISD) provided by the National Oceanic and Atmospheric Administration (NOAA). For each sampled line the days with conditions enabling the infection with Ha (RH >= 94%; Temp. 0-30°C) at the nearest station in the ISD divided by the total number of days with available data in every third year between 1986 – 2016 was calculated (Ha potential). **A**: The boxplots show the distribution of Ha potential for all lines from the respective population world-wide. Median and number of lines are shown above the boxplots. **B**: Equirectangular map of Europe showing the sampling locations of each line. The outline of the dots, representing one line each, indicate the respective population while the fill color indicates the Ha potential of the respective location (dark blue: low; yellow: high). The populations colors are ordered to enable maximum distinction between populations.

To asses if the low P_CHS_ in target regions of Ha-sRNAs in the coding region of At genes is the product of two sided evolution cumulating in balancing selection or adaptation in At cumulating in directional selection we estimated the *F_ST_* for all biallelic SNPs between the German and south Swedish populations with VCFtools (Danecek et al. 2011) after the method developed by Weir & Cockerham (1984) (Fig. 8). We estimated the *F_ST_* for 4,716,431 loci (median = 0.0116) in the genome of At (Fig. 9A). 204 (median = 0.0198) of these were located in target regions of the Ha-sRNA Set and 187, 211 and 194 (median = 0.0129, 0.0186, 0.0164) were located in target regions of the respective rsRNA Sets (Fig. 9B). Neither Set of *F_ST_* values differed significantly from one another and the genome.

**Figure 8:**
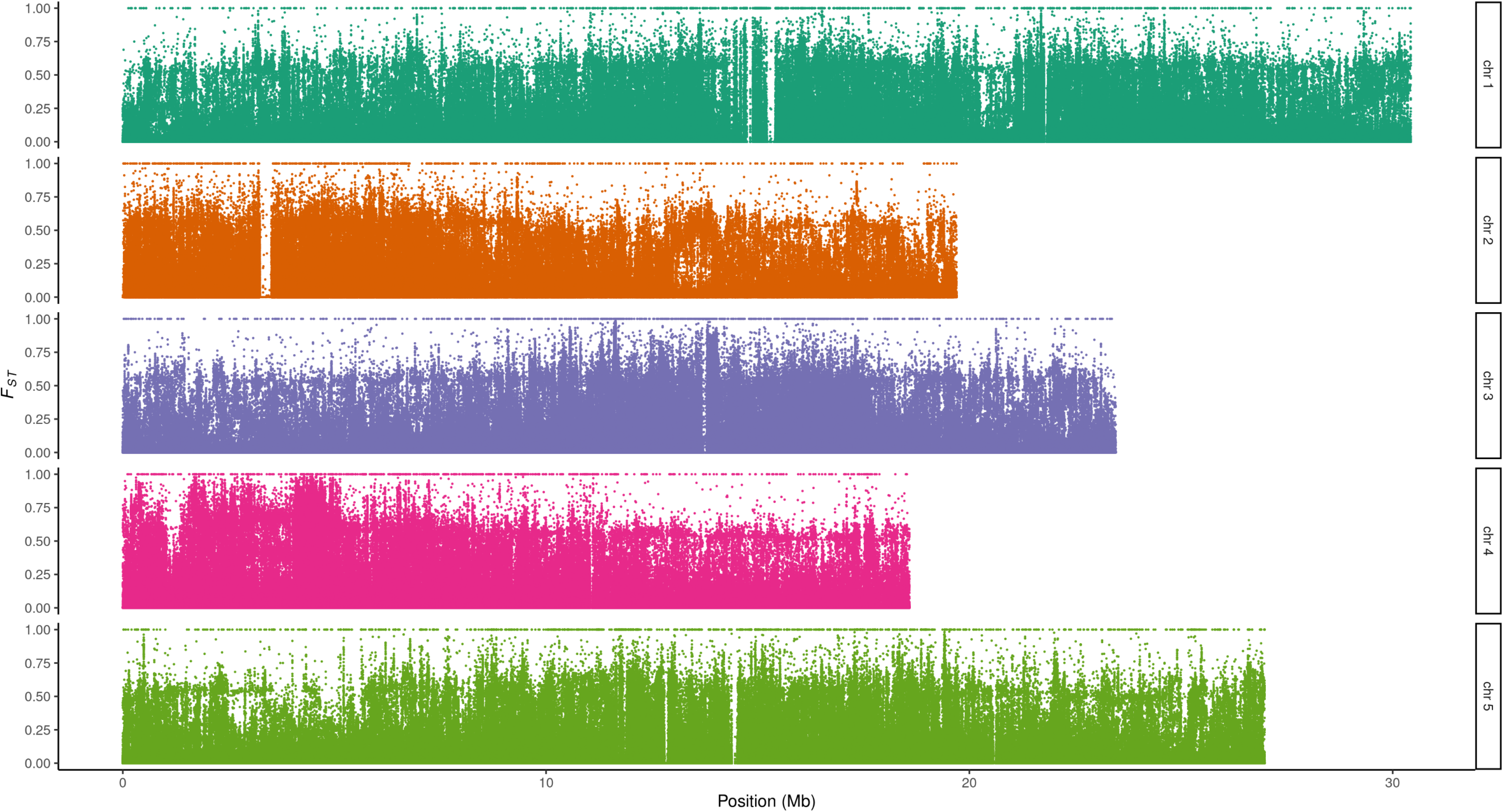
*F_ST_* values of each SNP shown for each chromosome in At. The *F_ST_* values between the German and south Swedish At populations were calculated with VCFtools. Dots indicate the *F_ST_* (y-axis) of a single locus. The positions in Mb are shown on the x-axis. Colors indicate chromosomes.

**Figure 9:**
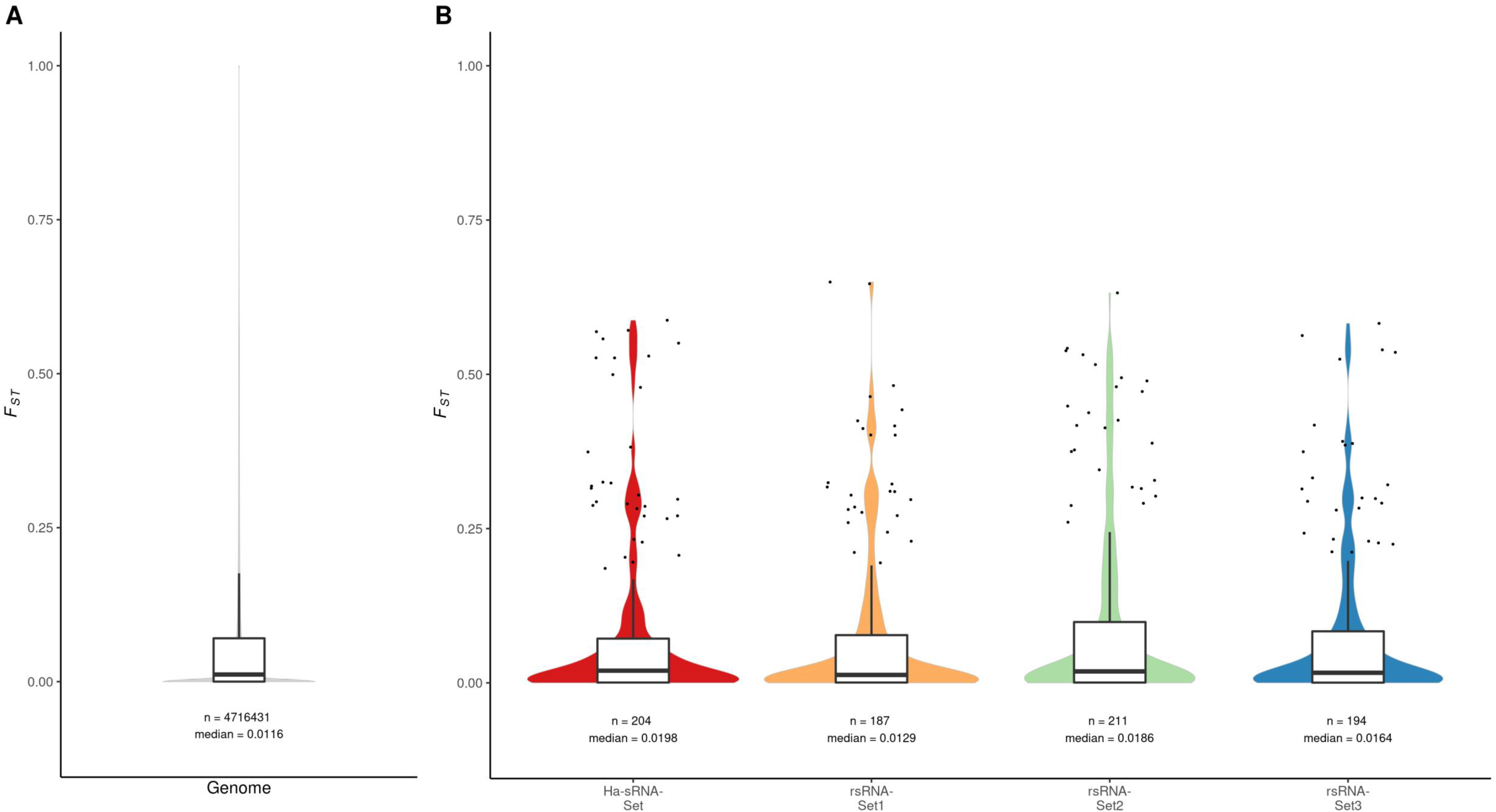
Distribution of *F_ST_* values in the genome and target regions of sRNA-sets. The *F_ST_* values between the German and south Swedish At populations were calculated with VCFtools. **A** shows the distribution of *F_ST_*s in the genome, **B** shows the distribution of *F_ST_* values in target regions of sRNA sets (red: Ha-sRNA Set, yellow: rsRNA Set1, green: rsRNA Set2, blue: rsRNA Set3). Boxplots and violin plots show the distribution of *F_ST_* values while outliers are shown as black dots. Number and median of *F_ST_* values in the respective set are shown below each boxplot. Outliers in **A** are not shown, due to being far too numerous.

While *F_ST_* Values in target regions of the Ha-sRNAs and respective rsRNA sets were not significantly different to the genome, *F_ST_* values and P_CHS_ values are negatively correlated exclusively for the Ha-sRNA Set (p = 0.03) (Fig. 10). This shows that target regions with an unlikely strong complementarity (low PCHS) are associated with higher *F_ST_* values, which suggests that these loci are under directional selection. These findings support that the unlikely high complementarity in many target regions of pathogen sRNAs is not the outcome of a two sided host-pathogen arms race which would lead to signs of balancing selection.

**Figure 10:**
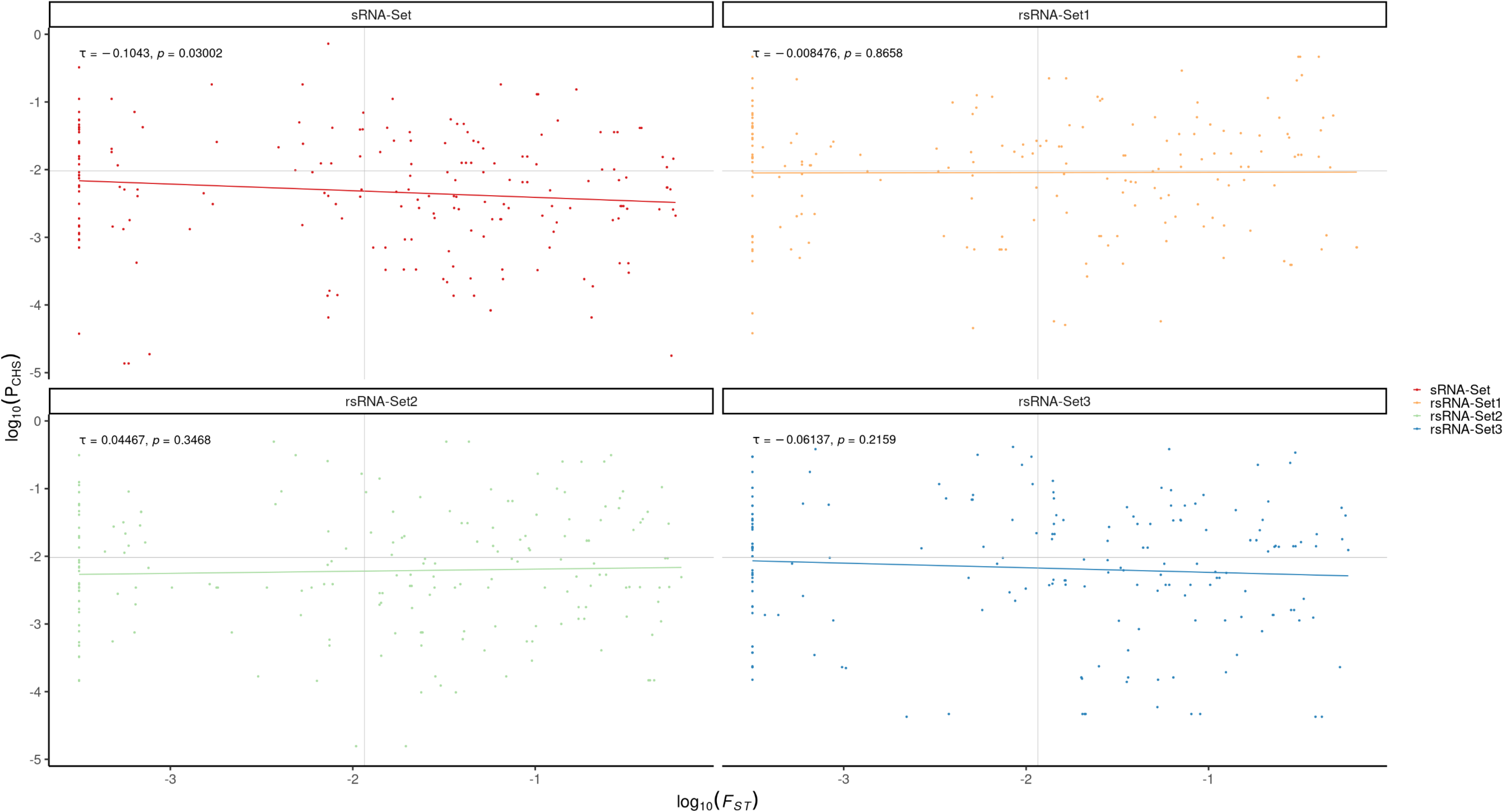
Correlation between *F_ST_* and P_CHS_ values in target regions of sRNA-sets. Calculated *F_ST_* values for each SNP (y-axis) are plotted against the P_CHS_ (x-axis) of the respective interaction. *F_ST_* and P_CHS_ are log_10_ transformed, while *F_ST_* = 0 is shown at 10^-3.5^. The p-value and τ of a Kendall rank correlation are shown in the top left corner of each plot. Within each plot a simple linear regression is shown. Grey vertical lines show the median *F_ST_* of the genomic background and grey horizontal lines indicate the median P_CHS_ of the rsRNA sets. Different colors indicate the respective sRNA set (red: Ha-sRNA Set, yellow: rsRNA Set1, green: rsRNA Set2, blue: rsRNA Set3).

Hypothetically, target regions of pathogen-derived sRNA can either be evolutionary constrained which would lead to no signs of natural selection and we would expect the P_CHS_ to be approximately the P_CHS_ in target regions of rsRNAs and *F_ST_* would be, due to the lack of allele differences between the populations undefined. On the other hand, if the target region of pathogen-derived sRNAs is unconstrained the effects on P_CHS_ and *F_ST_* would depend on two specific factors – fitness and evolutionary constraints of the pathogen-derived sRNA. If the pathogen-derived sRNA is evolutionary constrained, we would expect a directional selection of target regions (high *F_ST_*) and this target region would have an unlikely high (low P_CHS_) or low complementarity (high P_CHS_) depending whether if silencing of the target gene is beneficial or detrimental respectively in presence of the pathogen. If the pathogen-derived sRNA is evolutionary unconstrained we would expect a low *F_ST_* value indicating balancing selection while the interpretation of P_CHS_ in this case would be problematic (Fig. 11).

**Figure 11:**
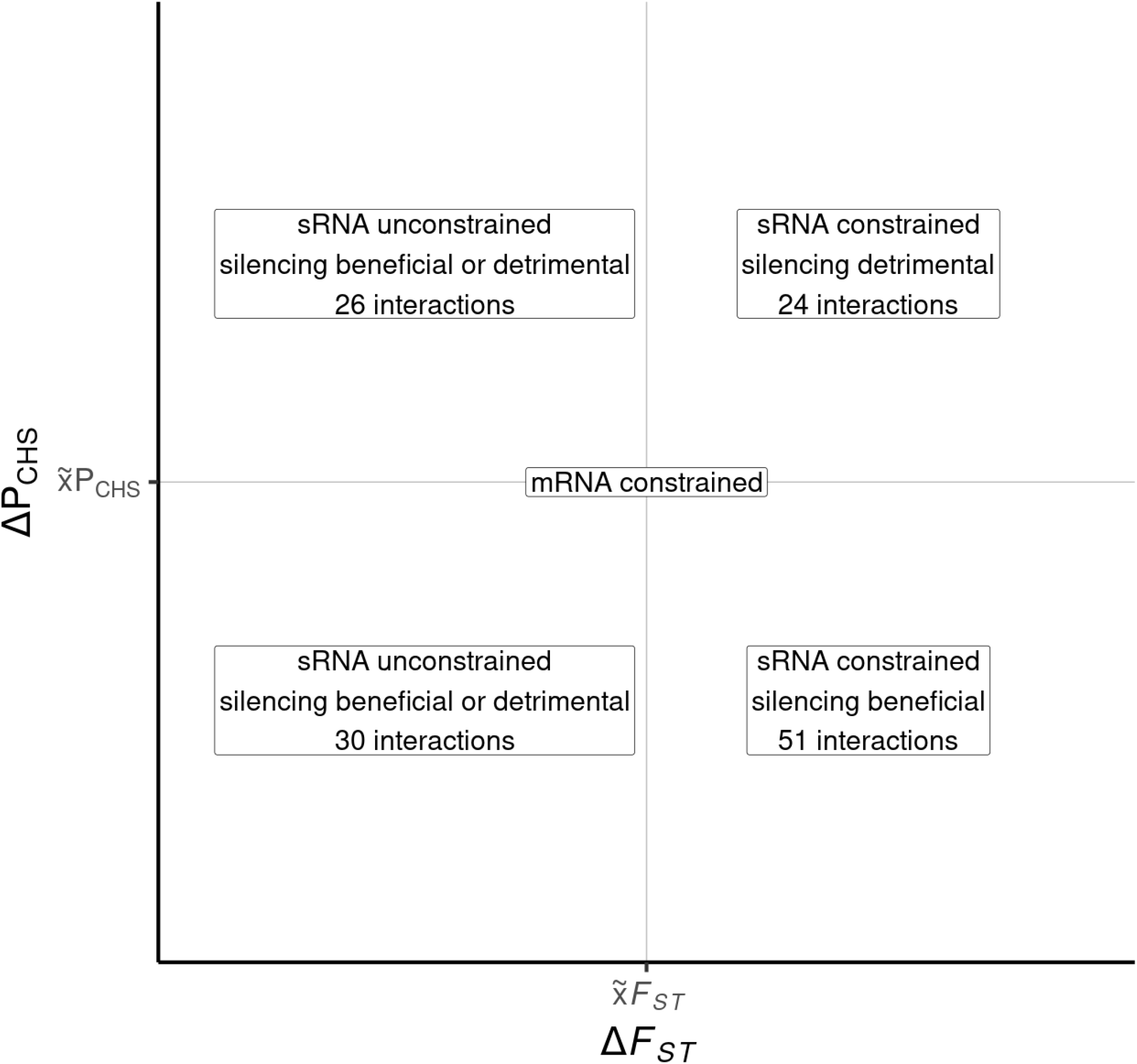
Hypothetical correlation between *F_ST_* and P_CHS_ in the context of evolutionary constraints and fitness effects and the number of interactions by the Ha-sRNA in each quadrant. While an evolutionary constrained mRNA will not show any deviations from the median *F_ST_* and P_CHS_ of the background, we expect to observe specific tendencies. If the ckRNAi exerting sRNA is able to evolve we will see a lower *F_ST_* due the presence of balancing selection (see Fig. 1). During balancing selection coevolution will prevent an unambiguous interpretation of P_CHS_ values (left quadrants). If ckRNAi exerting sRNAs are constrained and the target region of the mRNA is unconstrained and fitness effects are present we expect a directional selection, which will be expressed by an increase of *F_ST_* values (right quadrants). A beneficial effect of silencing would lead to an unexpectedly high complementarity (low PCHS; bottom right quadrant) and detrimental effects would lead to the opposite (high PCHS; top right quadrant). The number of predicted interactions by the Ha-sRNA Set with a respective P_CHS_ and a respective mean *F_ST_* in the target region are written inside the boxes.

From the 131 At-mRNAs coding regions targeted by Ha-sRNAs with available SNP data 75 show a higher *F_ST_* than the genome. Of these 51 show unexpectedly high complementarities to the respective sRNAs which is congruent with a beneficial effect of silencing for the plant, while only 24 interactions show the expected values of *F_ST_* and P_CHS_ congruent with the effector hypothesis. We conducted a set of binomial tests with the assumption that the probability of an interaction to fall in any quadrant separated by the genome-wide median *F_ST_* and the median P_CHS_ of rsRNA-sets was 25% for all interactions predicted for the Ha-sRNA Set and the three respective rsRNA sets. The 16 results were adjusted for multiple testing via the method of Benjamini & Yekutieli (2001) and only the high *F_ST_*/low P_CHS_ quadrant for the Ha-sRNA Set showed a significant increase (p= 0.029, BY adjusted).

A GO-term analysis of these 51 genes shows the enrichment of the jasmonate mediated signalling pathway, actin filament organization, establishment of localization, various developmental processes and alpha-amino acid metabolic processes (Fig. S4; Tab. S8). We ranked targeted genes with potential target sites of Ha-sRNAs and SNPs in the 1001 genomes dataset in their target regions by their mean *F_ST_* descending and *P_CHS_* ascending (Tab. 1). The gene AT1G52825 is exceptionally high ranked and is involved in mitochondrial cytochrome c oxidase (COX) assembly according to its GO annotation. Mitochondria are known to be essential for an extended ROS burst and COX is targeted in rice by the *Magnaporthe oryzae* effector Avr-Pita. This effector increases COX activity reducing the accumulation of reactive oxygen species and thereby preventing a hypersensitive response (Han et al. 2021). Our results suggest At evolved AT1G52825 to be silenced by Ha-sRNAs to disrupt COX assembly and enhancing hypersensitive responses.

**Tab. 1:**
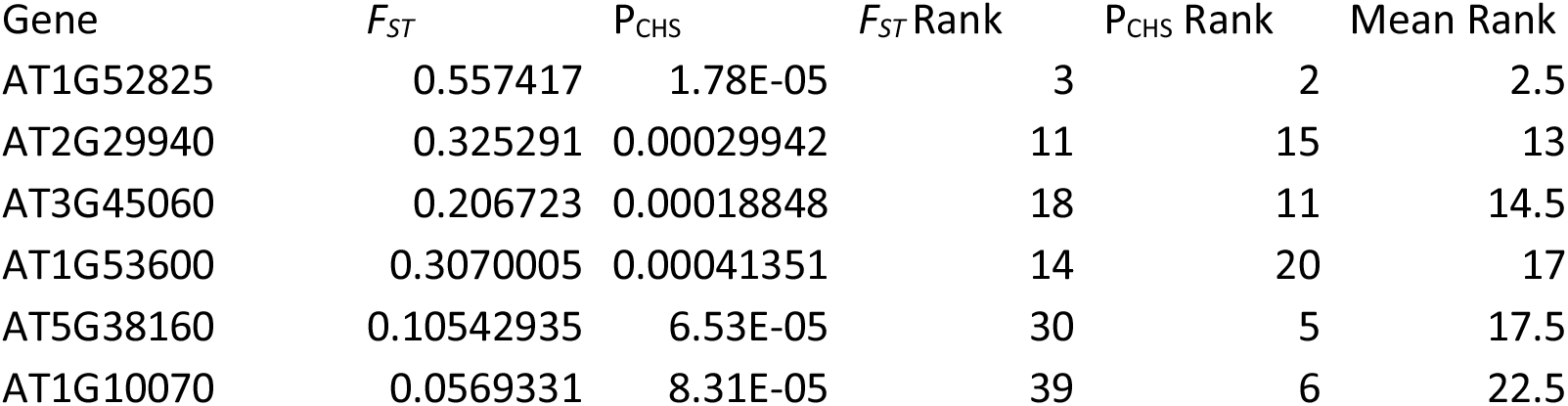
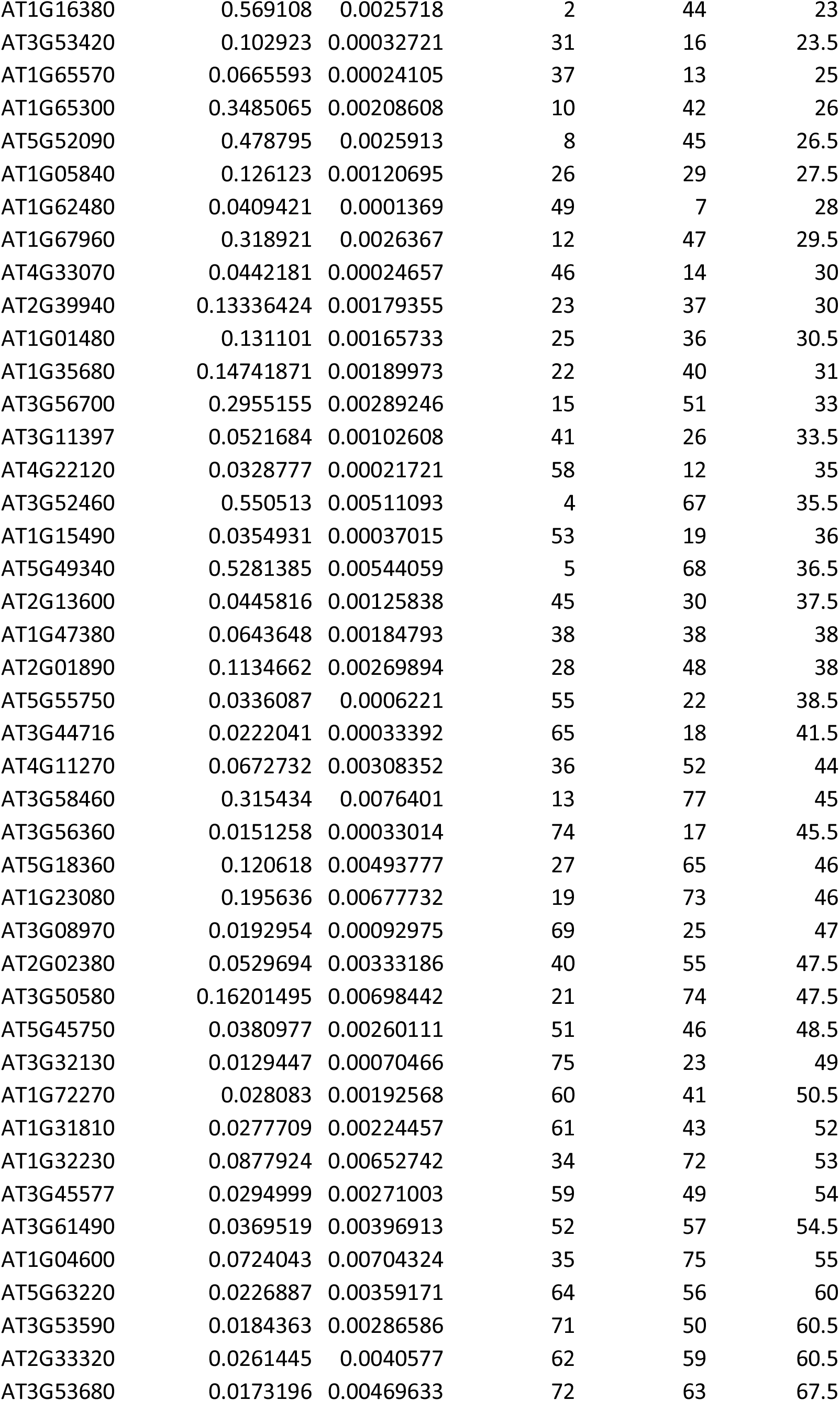
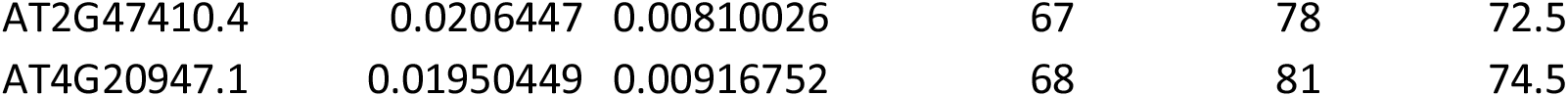
Ranking of At-genes by their *F_ST_* (descending) and P_CHS_ (ascending).

## Discussion

Unlike interaction prediction methods for protein effectors (Rao et al. 2014), the interactions of sRNA effectors with host genes can be predicted with less effort and higher reliability (Srivastava et al. 2014). In combination with the analysis of codon usage this untapped potential reveals a co-evolution within the hosts CDS. There is no reason why this co-evolution could not also take place outside the CDS, and our results therefore only give an incomplete picture.

Thus, according to current understanding, the natural occurrence of ckRNAi is harmful to the ingesting organism, but many organisms nevertheless ingest free exogenous dsRNA species (Qiang et al. 2021; Šečić and Kogel 2021). To solve these apparent conflicts, we propose an extended ckRNAi model in which organisms gather information about the microbiome of their habitat by taking up RNA from their environment to adapt to it, which mechanistically occurs via the RNAi pathway.

While the exact mechanism of sRNA transport between organisms during ckRNAi is still unresolved and debated (Nasfi & Kogel 2022), sRNAs traveling from pathogen to host and functions in silencing can be identified via AGO co-immunoprecipitation (Dunker et al. 2020). While this technique is applied to the question of pathogen-derived ckRNAi a sorting mechanism of sRNAs remains undescribed (Weiberg et al. 2013; Wang et al. 2016; Dunker et al. 2020). On the other hand, the sorting and delivery of protein effectors is governed by signal peptides, e.g. the conserved RxLR-motif in oomycetes (Whisson et al. 2007).

Contrastingly, specific plant-derived sRNAs are selected by RNA-binding proteins in the context of ckRNAi via extracellular vesicles (He et al. 2021). With the lack of evidence for a specific sorting mechanism for sRNA effectors by pathogens we postulate an, at least partially, unspecific transport of sRNAs from pathogen to host.

The majority of sRNA sequencing reads can be aligned to ribosomal (r)RNAs and transfer (t)RNAs and rRNA and tRNAs together account for 95% of RNA content in a typical eukaryotic cell, yet until recently were falsely dismissed from studies in the RNAi context and seen as mere contamination (Su et al. 2020; Chen et al. 2017). For several of these tRNA-derived fragments it was shown that these have functions in the ckRNAi context (Ren et al. 2019; Werner et al. 2020). Another abundant class of ckRNAi-exerting sRNA derive from transposable elements (Porquier et al. 2021). A common feature of these classes is the presence of multiple copies in the respective genomes and these genes fulfil sequence specific functions which constrains their evolution severely (Smit et al. 2007, Li et al. 2016; Zhang et al. 2020). Our extended view on ckRNAi aligns with these observations of unspecific transport of evolutionary constrained sRNAs.

This mechanism would benefit an organism by fine-tuning gene expression according to the presence of specific groups of pathogens, symbionts or commensalists. The extended model is not inconsistent with the current notion that certain pathogens “highjack” this mechanism (Weiberg et al. 2013), by delivering sRNA effectors. However, according to the extended model, this parasitic role of sRNAs is less prevalent and the uptake of environmental RNAs in general benefits organisms by providing a tool of microbiome perception.

Further research is necessary to identify the role of AT1G52825 and other highly ranked genes among the implicated beneficial targets of pathogen sRNAs in plant immunity.

## Supporting information

Supplemental Script

## Acknowledgments

This work was funded in the cooperative program of the Deutsche Forschungsgemeinschaft (DFG) FOR5116 to KHK.

## Competing interests

The authors declare that they have no competing interests.

## Availability of data and materials

The datasets supporting the conclusions of this article are available in the NCBI SRA repository, SRR11810702 (https://www.ncbi.nlm.nih.gov/sra/SRR11810702), SRR5852210 (https://www.ncbi.nlm.nih.gov/sra/SRR5852210) and SRR15248620 (https://www.ncbi.nlm.nih.gov/sra/SRR15248620).

## Supplementary

### Methods

#### sRNA-Dataset selection

Sequences for At-miRNAs and Gg-miRNAs were obtained from miRBase release 22.1 (miRBase.org; Kozomara et al. 2019). The whole dataset of miRNAs was read with the R package SeqinR v.3.6-1 (Charif & Lobry 2007) in the RStudio environment v.1.1.463 (RStudio Team 2020). Sequence names of miRNAs were selected with the R Base v.3.6.3 (R Core Team 2021) grep function and written to a different file for each organism.

For Ha-sRNAs the AtAGO1-co-IP datasets generated by Dunker et al. (2020) (SRR11810702, SRR5852210) were downloaded from the NCBI SRA with the sra toolkit function fastq-dump v.2.8.2 (Sherry et al. 2012) using the system2 package. Details for the Fg-sRNAs dataset from axenic culture are published in Werner et al. (2021).

#### Filtering of sRNAs

The Ha co-IP sequencing runs were treated similarly as described by Dunker et al. (2020). After downloading the already trimmed runs with fastq-dump the runs were transcribed to fasta with fastq_to_fasta and collapsed using the fastx_collapser from the fastx toolkit v0.0.14 (Gordon & Hannon 2010). The collapsed reads were first aligned to the At-TAIR10 (GCA_000001735) genome release (Lamesch et al. 2012) with the bowtie aligner v.1.2.1.1 (Langmead et al. 2009) with one allowed mismatch. Fg-sRNAs were not filtered to a plant genome due to their origin from axenic culture. Reads which did not align were kept and aligned to a Ha-mastergenome which consisted of the assemblies of the Ha-strains Emoy2, Cala2 and Noks1 (GCA_000173235.2, GCA_001414265.1, GCA_001414525.1) or the Fg-mastergenome which consisted of all 110 Fg genome assemblies from the NCBI database (see Table S7) and only perfect matches were kept. Remaining reads were further filtered with SeqinR and R Base for reads with at least a total sum of reads in both co-IP datasets of 200 a length between 21 and 24 nt and at least 1 RPM (reads per million) in each dataset for Ha, or 100 RPM for Fg.

#### sRNA clustering and generation of analogous random sRNA sets

sRNA-datasets were clustered with the CD-HIT v.4.8.1 (Fu et al. 2012) function cd-hit-est with a similarity threshold of 90%. The nucleotide frequency of clustered reads for position 1 (5’->3’), position 2-12 and the remaining nucleotides was calculated. These frequencies were used to generate three sets of analogous random sRNAs (rsRNAs) with the same number of sequences, the same frequency of nucleotides for each section and the same length distribution using SeqinR, the package stringr v.1.4.0 and the sample function from R Base with the relative frequencies as probabilities.

#### Preparation of At and barley coding sequences and calculation of codon usage indices

The CDS of the Araport11 annotation (Cheng et al. 2017) and *Hordeum vulgare IBSC PGSB v2* reference genome annotation (Mascher et al. 2017) were filtered and only CDS staring with a start codon (ATG) and ending with a stop codon (TAG, TAA, TGA) were retained. Additionally sequences were filtered for a length of a multiple of three nt’s using the function is.whole from the package sfsmisc v.1.1-7. Codon usage was calculated for the filtered sequences with the function uco from the SeqinR package.

#### Target prediction

For each set of sRNAs and its respective sets of analogous random sRNAs a target prediction was conducted with the TAPIR algorithm v.1.1 with a score cut-off of 6 (default=4) and a mfe-ratio of 0.6 (default=0.7) for At and a score cut-off of 8 and a mfe-ratio of 0.5 for barley according to the optimized parameters suggested by Srivastava et al. (2014) to obtain predicted likely interactions and those one mutation outside the default parameters.

#### P_CHS_ calculation

Target prediction results were read and saved to a R data frame and duplicated target regions of a specific sRNA in the same gene were removed with the duplicated function. The in frame CDS from the first sRNA overlapping codon was saved as a vector. The sRNA sequence was also saved in 3’-5’ direction in a character vector. Bulges in the mRNA or sRNA were accounted for by either removing the sRNA base overlapping the bulge in the sRNA sequence or by adding an unmatching character in the sRNA sequence for a bulge in the mRNA. After this preparation for each codon in the mRNA all synonymous codons were analyzed for their complementarity to the sRNA sequence with SeqinR and base functions and the probability of codons with 0, 1, 2 or 3 complementary bases to the sRNA were calculated based on the codon frequency calculated before. All possible 4 to the power of overlapping codons permutations were written to a data frame with the expand.grid function and the row sums were saved as a vector. The prior calculated probabilities of complementarities were used to replace the values in the permutations data frame and the prod function was applied to each row with the apply function giving the probabilities to each row sum of complementarities. The sum of probabilities for each row sum with i. the same complementarity as the actual interaction and ii. a higher complementarity were added to the data frame containing the target prediction results. The probabilities i. and ii. were added to calculate the P_CHS_ for each interaction. P_CHS_ values were plotted on log_10_-scale with the package ggplot2 v.3.3.2 (Wickham 2016) and ggpubr v.0.4.0 (Kassambara 2020). P-values were calculated via a Kruskal-Wallis test and adjusted for multiple testing (Benjamini & Yekutieli 2001) with the function compare_means. To calculate the 95% confidence intervals (CIs) the function t.test was applied with default parameters (two.sided, var.equal=F, paired=F, conf.level=0.95) on the approximately normally distributed log_10_(P_CHS_) values. These CIs were retransformed via 10^CI to obtain the relative difference between sRNA and rsRNA-sets.

#### Data acquisition

The full dataset of SNPs and indel was downloaded via the 1001 Genomes Data Center (https://1001genomes.org/data/GMI-MPI/releases/v3.1/1001genomes_snp-short-indel_with_tair10_only_ACGTN.vcf.gz) (Alonso-Blanco et al. 2016). Description of lines including Accession_ID, sampling location, names and population origin was downloaded via (https://tools.1001genomes.org/api/accessions.csv?query=SELECT%20*%20FROM%20tg_accessions%20ORDER%20BY%20id) and is from here on referred to as Accession_list. Subsequent analyses were conducted with R v.4.2.1 (R Core Team 2022) and RStudio v2021.09.0 (RStudio Team 2020). The weather data was downloaded with the rnoaa package v.1.3.8 (Chamberlain 2021). Stations in a radius of 250 km around each location from the Accession_list were searched and the closest station with data collection beginning before 1986-01-01 and ending after 2016-12-31 was selected via the function isd_stations_search. Weather data was obtained with the function isd for each previously selected station for every third year from 1986 to 2016. Obtained weather data was collected as a list of data frames.

#### Calculation of Ha potential

For each At-Accession the respective weather data was filtered for temperature measurements with optimal quality scores (1). Relative humidity (RH) was calculated for each measurement with the package humidity v.0.1.5 (Cai 2019) function “RH”. All measurements with a temperature above 30°C, below 0°C or a RH below 94% were removed to obtain a data frame only containing measurements with potential for Ha infection. The number of unique dates in the reduced data frame was divided by the number of unique dates in the complete data frame to calculate the relative amount of days with potential for Ha infection (Ha potential) for each accession.

#### Calculation of genome wide *F_ST_* values

The identifier of Accessions belonging to the German or south Swedish populations were extracted from the Accession_list and written to files. The 1001 genomes vcf-file was indexed and subsequently filtered with BCFtools v.1.11 (Danecek et al. 2021) (using htslib v. 1.11-4 (Bonfield et al. 2021)) index and view function respectively. The filtered file was used to estimate the *F_ST_* based on the method developed by Weir & Cockerham (1984) with the VCFtools v.0.1.16 (Danecek et al. 2011) function weir-fst-pop.

#### Selection of *F_ST_* values in target regions of sRNAs

For each interaction predicted in previous calculations using TAPIR (Bonnet et al. 2010) the genomic region was determined. To this end for each targeted transcripts CDS the chromosome_name, cds_start, cds_end, genomic_coding_start, genomic_coding_end and strand were obtained from https://plants.ensembl.org v.53 (Cunningham et al. 2022) via the package biomaRt v.2.50.3 (Durinck et al. 2005; Durinck et al. 2009). Additionally the VCFtools calculated *F_ST_* values were loaded into a data frame. Based on the start and end nt of the TAPIR results (from here on refered to as start/end) and the cds_start and cds_end values from biomaRt it was determined if the target region spans an exon-intron junction. If strand was 1 and no junction was present in the target region genomic start position and end position were calculated by start+genomic_coding_start-cds_start and end+genomic_coding_start-cds_start respectively, if strand was -1 without junction genomic start position and end position were calculated by genomic_coding_end-start+cds_start and genomic_coding_end-end+cds_start respectively. For the more complicated case of target regions spanning junctions genomic positions were saved to a list. For target regions in transcripts on the plus strand the start position was calculated via start+genomic_coding_start-cds_start and for the last exon genomic_coding_start and for the end positions respectively as genomic_coding_end or genomic_coding_start-cds_start+end. For target regions in transcripts on the minus strand positions were calculated as genomic_coding_end+cds_start-end, genomic_coding_start, genomic_coding_end and genomic_coding_start+cds_end-start respectively. *F_ST_* values were filtered from the data frame based on these regions. To ensure the selection of correct target regions genomic sequences from the At-TAIR10 assembly (GCA_000001735) were read with SeqinR and converted to uppercase strings. These strings were compared to the target region provided by the TAPIR results after removal of “-“ characters and a replacement of “U” with “T”. In the case of transcripts on the minus strands the sequence read from the At-TAIR10 assembly was before comparison reverse complemented with the function reverseComplement from the package Biostrings v.2.62.0 (Pagès et al. 2021).

#### Statistical analysis of *F_ST_* and P_CHS_ values

Due to the estimation method developed by Weir & Cockerham (1984) some *F_ST_* values were calculated as slightly below 0 and were changed to 0 in the data frame before statistical analyses. Significant differences between genome-wide *F_ST_* and *F_ST_* in target regions of sRNA-sets comparisons were conducted in the same manner as the comparison of P_CHS_ values via a Kruskal-Wallis test and adjusted for multiple testing (Benjamini & Yekutieli 2001) with the function compare_means from the package ggpubr v.0.4.0 (Kassambara 2020). No significant differences were detected. Kendall correlations between *F_ST_* and P_CHS_ were calculated and plotted with the ggpubr function stat_cor (two.sided). Binomial tests for the abundance of gene target region interactions on the basis of quadrants separated by genome-wide median *F_ST_* and the median P_CHS_ of rsRNA-sets was conducted with the function binom.test (hypothesised probability = 0.25; two.sided) and adjusted for multiple testing with the function p.adjust (method=BY) from the package stats.

Custom code is provided in Additional File 1 (supplementary_script.r.txt)

---

**Figure S1:**
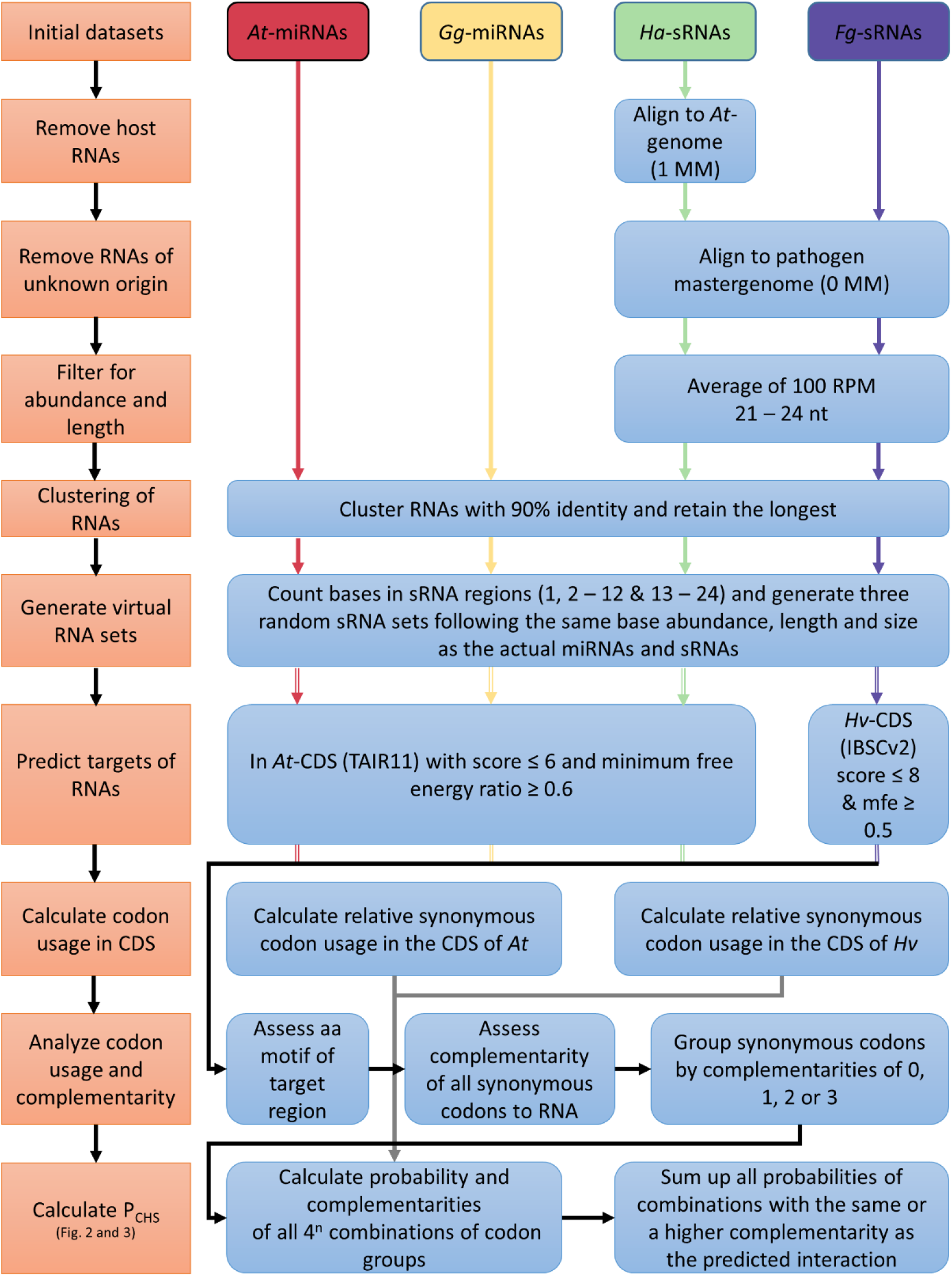
Workflow of the computational analysis of codon usage in RNAi targeted mRNA regions. The left column (orange) gives an overview of the overall workflow. More detailed steps are depicted in blue. The colored arrows (red: At*, Arabidopsis thaliana*; yellow: Gg, *Gorilla gorilla*; green: Ha, *Hyaloperonospora arabidopsidis*; purple: Fg, *Fusarium graminearum*) indicate each set of small RNAs by organism of origin and visualize the detailed steps applied to each set, while black and gray arrows apply to all sets. In 4^n^, n refers to the number of codons covered fully or partially by the respective sRNA, which leads to a maximum of 4^9^ combinations for each interaction, in the case of a not in frame 24 nt long sRNA covering 7 codons completely and 2 partly. For further details, please refer to the supplementary methods.

**Figure S2:**
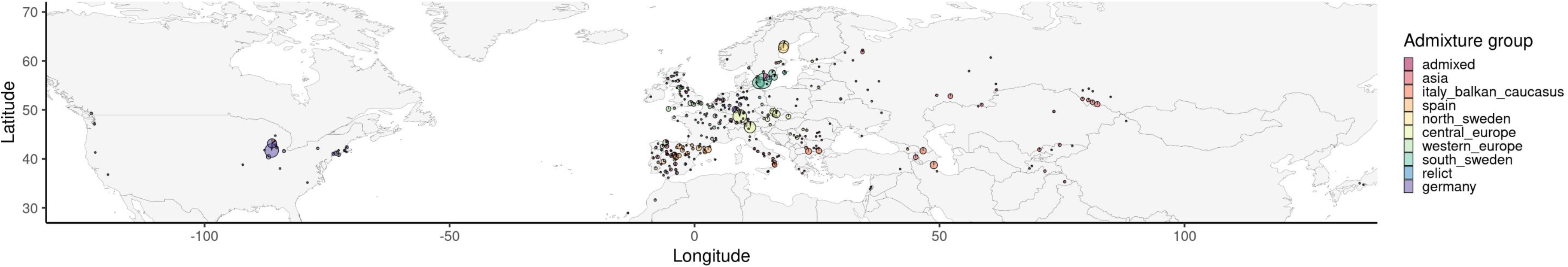
Weather stations used for the climate estimates of At populations (world). This equirectangular map shows locations of all weather stations used to estimate the climatic conditions for each population. Each pie chart represents one station. Size indicates the number of lines represented by the station and colors indicate the respective population of lines represented.

**Figure S3:**
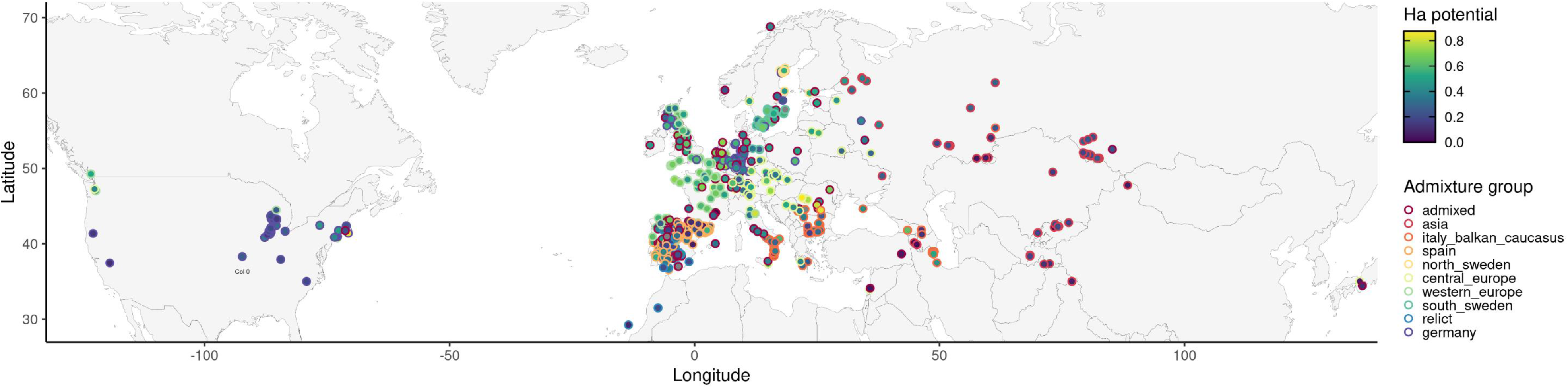
Estimated potential for Ha infections (world map). Equirectangular map showing the potential for Ha infections, estimated by the analysis of weather data from the Integrated Surface Database (ISD) provided by the National Oceanic and Atmospheric Administration (NOAA). For each sampled line the days with conditions enabling the infection with Ha (RH >= 94%; Temp. 0-30°C) at the nearest station in the ISD divided by the total number of days with available data in every third year between 1986 – 2016 was calculated (Ha potential). Each dot shows the sampling location of one line. The outline of the dots indicates the respective population while the fill color indicates the Ha potential of the respective location (dark blue: low; yellow: high). The populations colors are ordered to enable maximum distinction between populations.

**Figure S4:**
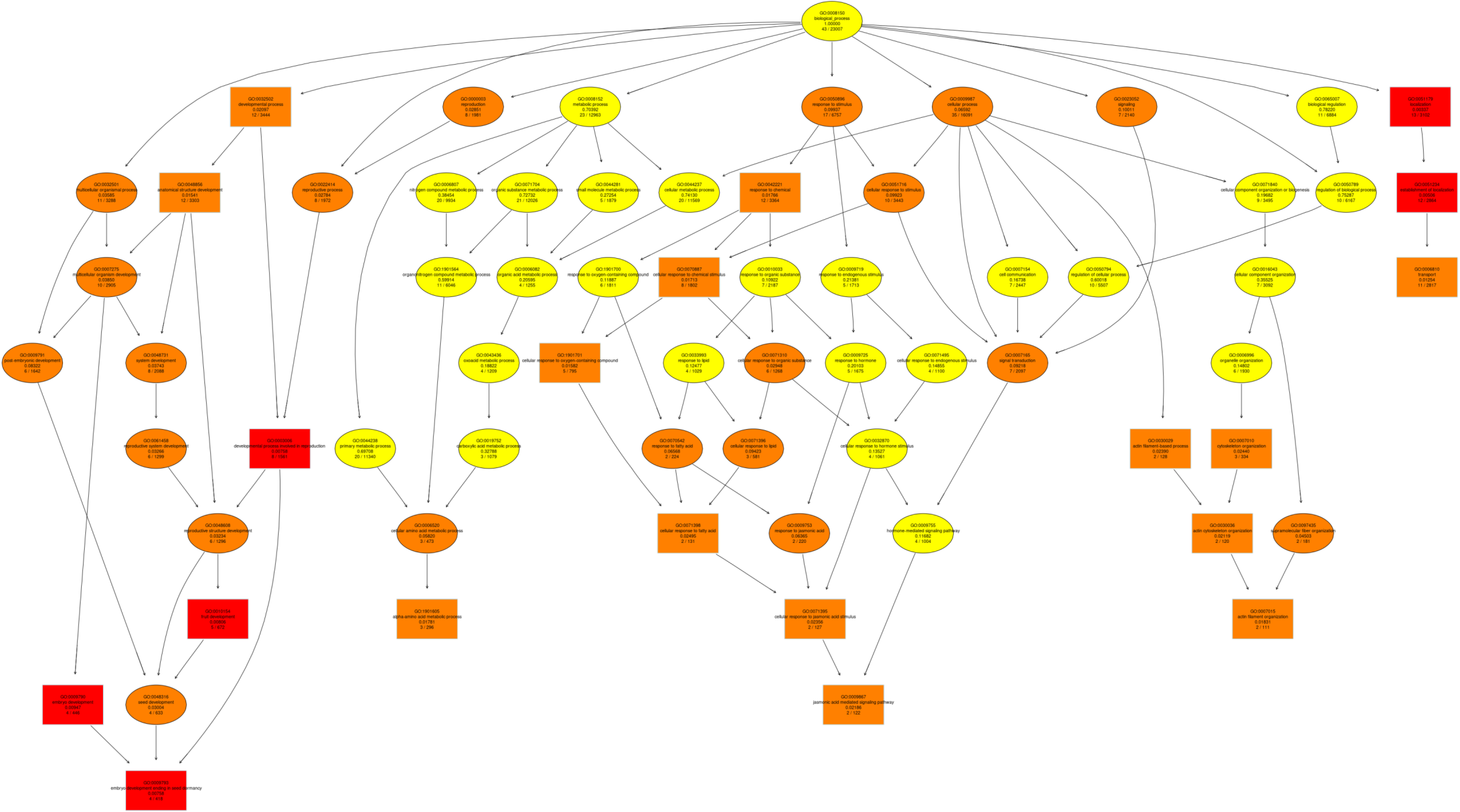
Hierarchical structure of enriched GO-terms (p<0.025) in the target genes of Ha-sRNAs showing low P_CHS_ and high *F_ST_* values. The graph shows the significantly enriched GO-terms and parent terms of target genes of Ha-sRNAs showing low P_CHS_ and high *F_ST_* values. Colors indicate significance from dark red (most significant) to yellow (less significant). The GO-identifier, GO-term name, the p-value (unadjusted), the number of annotated and the number of targeted genes are shown within boxes. To obtain the most relevant results enrichment analysis was restricted to terms with 100 or more annotations in the At gene set.

**Tab. S1:**
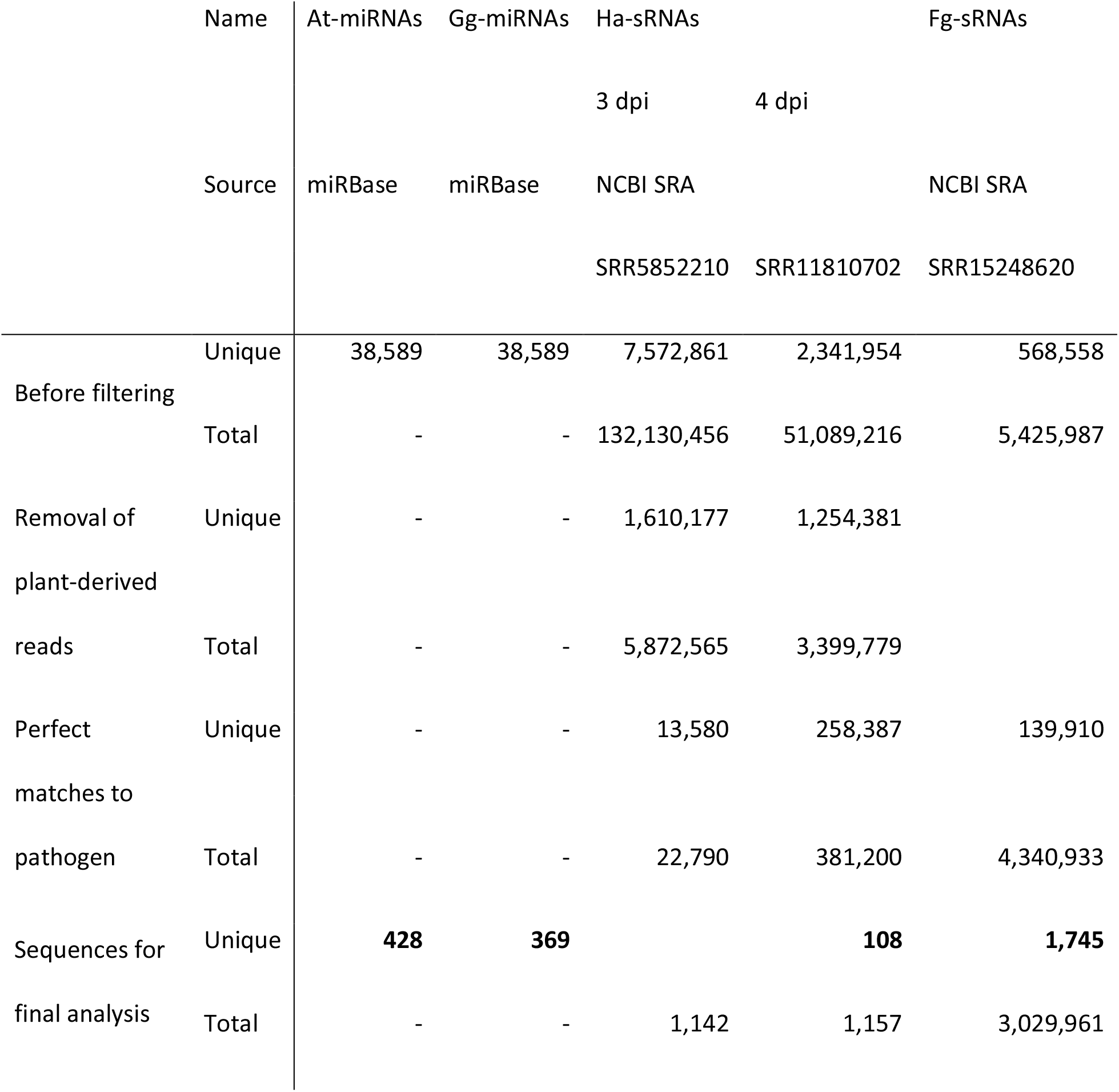
Summary of sRNA datasets This table gives the number of reads or sequences (total and unique) before and after each filtering step with the number of sequences kept for further analysis in bold.

**Tab. S2:** Summary of sRNA clustering This table shows the number of sequences after the filtering (shown in Tab. S1) and the reduction of reads or sequences after clustering of similar RNAs.

| RNA set | At-miRNAs | Gg-miRNAs | Ha-sRNAs | Fg-sRNAs |
| --- | --- | --- | --- | --- |
| No of sequences | 428 | 369 | 108 | 1,745 |
| No of clusters | **335** | **351** | **66** | **457** |
| Mean cluster size | 1.278 | 1.051 | 1.636 | 3.818 |

**Tab. S3:** Nucleotide composition of sRNA sets in different sections This table gives the base composition in the clustered sRNA-sets used for the generation of the random analogous RNA-sets.

| RNA set | At-miRNAs | Gg-miRNAs | Ha-sRNAs | Fg-sRNAs |
| --- | --- | --- | --- | --- |
| 1st nt |  |  |  |  |
| A | 24.8% | 30.5% | 30.3% | 23.9% |
| C | 9.3% | 14.2% | 21.2% | 23.2% |
| G | 10.7% | 8.8% | 22.7% | 20.4% |
| U | 55.2% | 46.4% | 25.8% | 32.6% |
| Seed (2nd-12th nt) |  |  |  |  |
| A | 26.6% | 26.1% | 28.5% | 24.1% |
| C | 18.2% | 23.0% | 19.4% | 21.1% |
| G | 26.1% | 26.6% | 26.7% | 29.2% |
| U | 29.1% | 24.3% | 25.3% | 25.6% |
| End (13th-last nt) |  |  |  |  |
| A | 27.3% | 23.0% | 24.0% | 23.2% |
| C | 19.4% | 21.0% | 18.9% | 23.4% |
| G | 24.2% | 27.9% | 30.5% | 28.4% |
| U | 29.1% | 28.1% | 26.7% | 25.1% |
| Total |  |  |  |  |
| A | 26.8% | 25.0% | 26.5% | 23.6% |
| C | 18.3% | 21.7% | 19.2% | 22.3% |
| G | 24.5% | 26.3% | 28.3% | 28.4% |
| U | 30.3% | 27.0% | 26.0% | 25.7% |

**Tab. S4:** Occurrences and relative frequencies of synonymous codons in the Araport11 annotation

| Amino Acid | Codon | Occurrences | Relative Frequency |
| --- | --- | --- | --- |
| Lysine | AAA | 659,375 | 49.59% |
|  | AAG | 670,179 | 50.41% |
| Asparagine | AAC | 418,657 | 45.77% |
|  | AAT | 496,098 | 54.23% |
| Threonine | ACA | 342,305 | 32.50% |
|  | ACC | 199,360 | 18.93% |
|  | ACG | 147,692 | 14.02% |
|  | ACT | 363,987 | 34.56% |
| Arginine | AGA | 408,475 | 36.28% |
|  | AGG | 231,221 | 20.53% |
|  | CGA | 131,881 | 11.71% |
|  | CGC | 76,766 | 6.82% |
|  | CGG | 100,347 | 8.91% |
|  | CGT | 177,310 | 15.75% |
| Serine | AGC | 242,457 | 12.64% |
|  | AGT | 315,625 | 16.46% |
|  | TCA | 408,100 | 21.28% |
|  | TCC | 226,420 | 11.81% |
|  | TCG | 184,065 | 9.60% |
|  | TCT | 540,891 | 28.21% |
| Isoleucine | ATA | 284,518 | 25.85% |
|  | ATC | 358,653 | 32.58% |
|  | ATT | 457,619 | 41.57% |
| Methionine | ATG | 509,709 | 100.00% |
| Glutamine | CAA | 420,938 | 56.37% |
|  | CAG | 325,843 | 43.63% |
| Histidine | CAC | 173,590 | 36.58% |
|  | CAT | 300,965 | 63.42% |
| Proline | CCA | 340,804 | 34.32% |
|  | CCC | 106,564 | 10.73% |
|  | CCG | 158,588 | 15.97% |
|  | CCT | 387,002 | 38.97% |
| Leucine | CTA | 215,166 | 10.82% |
|  | CTC | 311,189 | 15.65% |
|  | CTG | 219,166 | 11.02% |
|  | CTT | 513,875 | 25.84% |
|  | TTA | 280,005 | 14.08% |
|  | TTG | 449,182 | 22.59% |
| Glutamic Acid | GAA | 753,248 | 53.03% |
|  | GAG | 667,114 | 46.97% |
| Aspartic Acid | GAC | 345,908 | 30.60% |
|  | GAT | 784,565 | 69.40% |
| Alanine | GCA | 377,962 | 28.88% |
|  | GCC | 195,418 | 14.93% |
|  | GCG | 167,370 | 12.79% |
|  | GCT | 567,989 | 43.40% |
| Glycine | GGA | 486,845 | 36.76% |
|  | GGC | 183,090 | 13.82% |
|  | GGG | 211,801 | 15.99% |
|  | GGT | 442,665 | 33.42% |
| Valine | GTA | 220,305 | 15.96% |
|  | GTC | 248,294 | 17.99% |
|  | GTG | 349,466 | 25.32% |
|  | GTT | 562,053 | 40.72% |
| Stop codon | TAA | 17,577 | 36.44% |
|  | TAG | 9,898 | 20.52% |
|  | TGA | 20,757 | 43.04% |
| Tyrosine | TAC | 265,603 | 45.56% |
|  | TAT | 317,324 | 54.44% |
| Cysteine | TGC | 153,423 | 40.25% |
|  | TGT | 227,727 | 59.75% |
| Tryptophan | TGG | 254,759 | 100.00% |
| Phenylalanine | TTC | 403,565 | 46.05% |
|  | TTT | 472,797 | 53.95% |

**Tab. S5:** Occurrences and relative frequencies of synonymous codons in the barley IBSCv2 annotation

| Amino Acid | Codon | Occurrences | Relative Frequency |
| --- | --- | --- | --- |
| Lysine | AAA | 931,360 | 37.85% |
|  | AAG | 1,529,125 | 62.15% |
| Asparagine | AAC | 886,774 | 49.86% |
|  | AAT | 891,820 | 50.14% |
| Threonine | ACA | 692,223 | 30.38% |
|  | ACC | 607,873 | 26.68% |
|  | ACG | 392,499 | 17.23% |
|  | ACT | 585,985 | 25.72% |
| Arginine | AGA | 577,809 | 21.47% |
|  | AGG | 704,744 | 26.19% |
|  | CGA | 235,490 | 8.75% |
|  | CGC | 443,708 | 16.49% |
|  | CGG | 413,389 | 15.36% |
|  | CGT | 315,934 | 11.74% |
| Serine | AGC | 755,547 | 18.90% |
|  | AGT | 550,365 | 13.77% |
|  | TCA | 783,348 | 19.59% |
|  | TCC | 681,459 | 17.04% |
|  | TCG | 446,272 | 11.16% |
|  | TCT | 781,049 | 19.54% |
| Isoleucine | ATA | 567,069 | 25.17% |
|  | ATC | 870,434 | 38.63% |
|  | ATT | 815,806 | 36.20% |
| Methionine | ATG | 1,209,257 | 100.00% |
| Glutamine | CAA | 716,645 | 41.25% |
|  | CAG | 1,020,702 | 58.75% |
| Histidine | CAC | 549,041 | 46.03% |
|  | CAT | 643,733 | 53.97% |
| Proline | CCA | 718,205 | 30.34% |
|  | CCC | 457,329 | 19.32% |
|  | CCG | 515,556 | 21.78% |
|  | CCT | 675,991 | 28.56% |
| Leucine | CTA | 434,057 | 9.50% |
|  | CTC | 965,930 | 21.14% |
|  | CTG | 1,045,193 | 22.88% |
|  | CTT | 918,147 | 20.10% |
|  | TTA | 390,168 | 8.54% |
|  | TTG | 815,116 | 17.84% |
| Glutamic Acid | GAA | 1,203,764 | 42.61% |
|  | GAG | 1,621,566 | 57.39% |
| Aspartic Acid | GAC | 1,136,016 | 45.29% |
|  | GAT | 1,372,440 | 54.71% |
| Alanine | GCA | 959,211 | 26.26% |
|  | GCC | 1,004,024 | 27.49% |
|  | GCG | 684,449 | 18.74% |
|  | GCT | 1,004,609 | 27.51% |
| Glycine | GGA | 747,093 | 23.57% |
|  | GGC | 1,002,984 | 31.65% |
|  | GGG | 666,870 | 21.04% |
|  | GGT | 752,150 | 23.73% |
| Valine | GTA | 417,867 | 13.49% |
|  | GTC | 777,749 | 25.11% |
|  | GTG | 1,004,162 | 32.42% |
|  | GTT | 897,389 | 28.97% |
| Stop codon | TAA | 30,806 | 24.20% |
|  | TAG | 36,006 | 28.29% |
|  | TGA | 60,444 | 47.49% |
| Tyrosine | TAC | 691,781 | 54.49% |
|  | TAT | 577,788 | 45.51% |
| Cysteine | TGC | 540,108 | 59.38% |
|  | TGT | 369,471 | 40.62% |
| Tryptophan | TGG | 593,276 | 100.00% |
| Phenylalanine | TTC | 997,261 | 55.90% |
|  | TTT | 786,642 | 44.10% |

**Tab. S6:** Summary of target prediction results This table gives an overview of the target prediction results and the subsequent filtering with the last column giving the remaining interactions for statistics and plotting

| Organism | RNA set | Interactions | sRNAs<br>targeting | Targeted<br>genes | Unique<br>gene-sRNA<br>interactions |
| --- | --- | --- | --- | --- | --- |
| At | actual | 14,524 | 335 | 6,318 | 8,124 |
|  | analogous 1 | 8,936 | 333 | 4,195 | 4,728 |
|  | analogous 2 | 9,069 | 329 | 4,279 | 4,866 |
|  | analogous 3 | 8,586 | 333 | 4,156 | 4,700 |
| Gg | actual | 8,762 | 348 | 4,023 | 4,750 |
|  | analogous 1 | 6,767 | 338 | 3,323 | 3,650 |
|  | analogous 2 | 7,244 | 342 | 3,538 | 3,928 |
|  | analogous 3 | 6,980 | 339 | 3,450 | 3,822 |
| Ha | actual | 783 | 59 | 421 | 438 |
|  | analogous 1 | 739 | 57 | 410 | 418 |
|  | analogous 2 | 645 | 59 | 392 | 396 |
|  | analogous 3 | 701 | 54 | 366 | 372 |
| Fg | actual | 12,649 | 452 | 2,322 | 3,155 |
|  | analogous 1 | 11,183 | 447 | 2,458 | 2,686 |
|  | analogous 2 | 12,038 | 448 | 2,546 | 2,793 |
|  | analogous 3 | 11,676 | 449 | 2,476 | 2,739 |

**Tab. S7:**
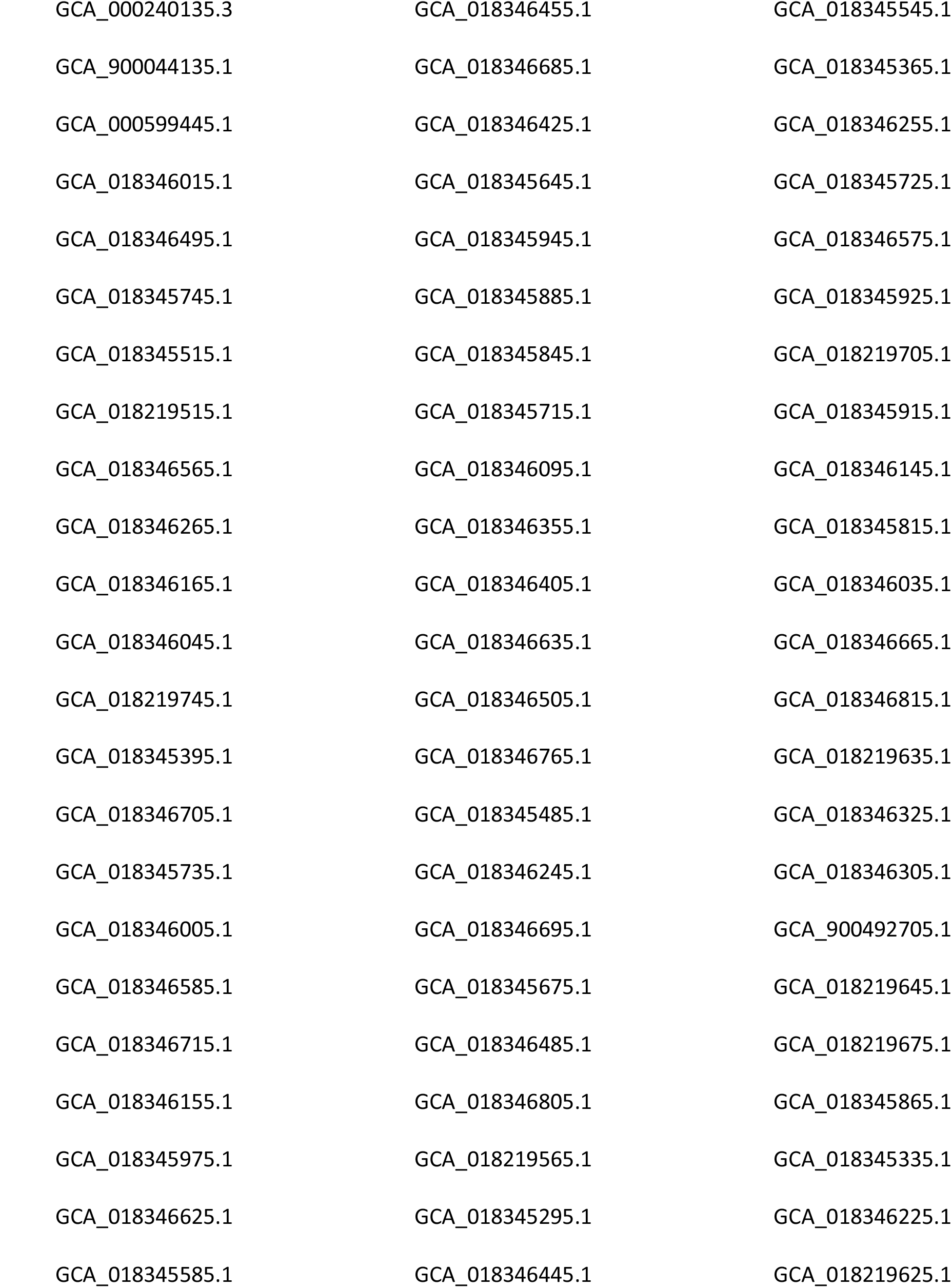
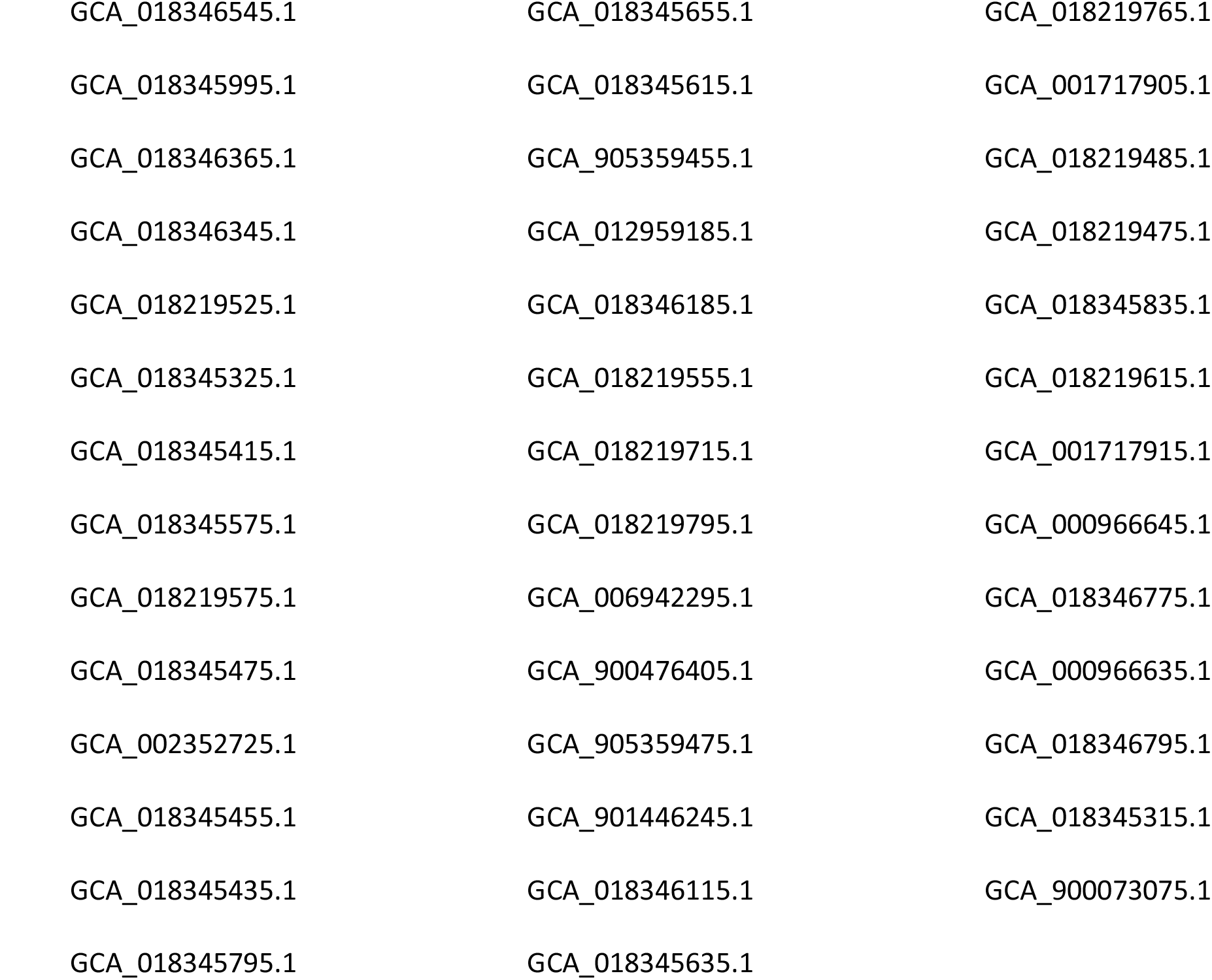
List of Fg-assembly’s accessions used for alignment of Fg-reads.

**Tab. S8:** Significantly enriched GO terms in the set of 51 genes showing a low P_CHS_ and high *F_ST_*.

| GO.ID | Term | Annotated | Significant | Expected | p-value |
| --- | --- | --- | --- | --- | --- |
| GO:0051179 | localization | 3102 | 13 | 5.8 | 0.0034 |
| GO:0051234 | establishment of localization | 2864 | 12 | 5.35 | 0.0051 |
| GO:0009793 | embryo development ending in seed dormancy | 418 | 4 | 0.78 | 0.0076 |
| GO:0003006 | developmental process involved in reproduction | 1561 | 8 | 2.92 | 0.0076 |
| GO:0010154 | fruit development | 672 | 5 | 1.26 | 0.0081 |
| GO:0009790 | embryo development | 446 | 4 | 0.83 | 0.0095 |
| GO:0006810 | transport | 2817 | 11 | 5.26 | 0.0125 |
| GO:0048856 | anatomical structure development | 3303 | 12 | 6.17 | 0.0154 |
| GO:1901701 | cellular response to oxygen-containing compound | 795 | 5 | 1.49 | 0.0158 |
| GO:0070887 | cellular response to chemical stimulus | 1802 | 8 | 3.37 | 0.0171 |
| GO:0042221 | response to chemical | 3364 | 12 | 6.29 | 0.0177 |
| GO:1901605 | alpha-amino acid metabolic process | 296 | 3 | 0.55 | 0.0178 |
| GO:0007015 | actin filament organization | 111 | 2 | 0.21 | 0.0183 |
| GO:0032502 | developmental process | 3444 | 12 | 6.44 | 0.021 |
| GO:0030036 | actin cytoskeleton organization | 120 | 2 | 0.22 | 0.0212 |
| GO:0009867 | jasmonic acid mediated signaling pathway | 122 | 2 | 0.23 | 0.0219 |
| GO:0071395 | cellular response to jasmonic acid stimulus | 127 | 2 | 0.24 | 0.0236 |
| GO:0030029 | actin filament-based process | 128 | 2 | 0.24 | 0.0239 |
| GO:0007010 | cytoskeleton organization | 334 | 3 | 0.62 | 0.0244 |
| GO:0071398 | cellular response to fatty acid | 131 | 2 | 0.24 | 0.025 |
| GO:1901698 | response to nitrogen compound | 340 | 3 | 0.64 | 0.0255 |
| GO:0022414 | reproductive process | 1972 | 8 | 3.69 | 0.0278 |
| GO:0048364 | root development | 618 | 4 | 1.16 | 0.0278 |
| GO:0022622 | root system development | 619 | 4 | 1.16 | 0.028 |
| GO:0000003 | reproduction | 1981 | 8 | 3.7 | 0.0285 |
| GO:0071310 | cellular response to organic substance | 1268 | 6 | 2.37 | 0.0295 |
| GO:0048316 | seed development | 633 | 4 | 1.18 | 0.03 |
| GO:0048608 | reproductive structure development | 1296 | 6 | 2.42 | 0.0323 |
| GO:0061458 | reproductive system development | 1299 | 6 | 2.43 | 0.0327 |
| GO:0032501 | multicellular organismal process | 3288 | 11 | 6.15 | 0.0358 |
| GO:0055085 | transmembrane transport | 1340 | 6 | 2.5 | 0.0372 |
| GO:0048731 | system development | 2088 | 8 | 3.9 | 0.0374 |
| GO:0033036 | macromolecule localization | 1342 | 6 | 2.51 | 0.0374 |
| GO:0007275 | multicellular organism development | 2905 | 10 | 5.43 | 0.0385 |
| GO:0016311 | dephosphorylation | 401 | 3 | 0.75 | 0.0388 |
| GO:0090502 | RNA phosphodiester bond hydrolysis, endonucleolytic | 170 | 2 | 0.32 | 0.0402 |
| GO:0009653 | anatomical structure morphogenesis | 1034 | 5 | 1.93 | 0.0426 |
| GO:0006869 | lipid transport | 176 | 2 | 0.33 | 0.0428 |
| GO:0097435 | supramolecular fiber organization | 181 | 2 | 0.34 | 0.045 |
| GO:0034613 | cellular protein localization | 734 | 4 | 1.37 | 0.0475 |
| GO:1901607 | alpha-amino acid biosynthetic process | 191 | 2 | 0.36 | 0.0496 |

## Notes

### Competing Interest Statement

The authors have declared no competing interest.

