## Supplemental Script for "Analysis of codon usage and allele frequencies reveal the double-edged nature of cross-kingdom RNAi"

#Generation of rsRNAs analogous to an actual set
library(seqinr)
library(stringr)
YOUR_Set<-""   #include folder

sRNAset<-read.fasta(file=YOUR_Set,as.string=TRUE)

sRNAset<-unlist(sRNAset,recursive=F)
sRNAset<-as.data.frame(sRNAset)
names(sRNAset)<-c("Seq")
sRNAset$Seq<-as.vector(sRNAset$Seq)
sRNAset$seed<-substr(sRNAset$Seq,2,12)
sRNAset$first<-substr(sRNAset$Seq,1,1)
sRNAset$end<-substr(sRNAset$Seq,13,30)
Af<-str_count(sRNAset$first,"a")
wfr<-table(nchar(as.vector(sRNAset$Seq)))
wfr<-as.data.frame(wfr)
basefreq<-data.frame(region=NA,a=NA,c=NA,g=NA,t=NA)
for(i in names(sRNAset)){
  a<-sum(str_count(sRNAset[[i]],"a"))
  c<-sum(str_count(sRNAset[[i]],"c"))
  g<-sum(str_count(sRNAset[[i]],"g"))
  t<-sum(str_count(sRNAset[[i]],"u"))
  if (t==0){t<-sum(str_count(sRNAset[[i]],"t"))}
  s<-sum(c(a,c,g,t))
  
  basefreq<-rbind(basefreq,(c(i,(a/s),(c/s),(g/s),(t/s))))
  
}
wfr$Freq<-as.numeric(as.character(wfr$Freq))
wfr$Var1<-as.numeric(as.character(wfr$Var1))
for (iii in c("Set1","Set2","Set3")){
  for (ii in 1:length(wfr$Freq)){
    for (i in 1:wfr$Freq[ii]) {
      line_header = paste(i,wfr$Var1[ii], sep="_")
      DNA = c ("A","T","C","G") 
      DNA_sequence = paste(paste(sample(DNA, 
                                        1, 
                                        replace=TRUE, 
                                        prob=c(basefreq$a[4],basefreq$t[4]
                                               ,basefreq$c[4],basefreq$g[4])),collapse =""),
                           paste(sample(DNA, 
                                        11, 
                                        replace=TRUE, 
                                        prob=c(basefreq$a[3],basefreq$t[3]
                                               ,basefreq$c[3],basefreq$g[3])),collapse =""),
                           paste(sample(DNA, 
                                        (wfr$Var1[ii]-12), 
                                        replace=TRUE, 
                                        prob=c(basefreq$a[5],basefreq$t[5]
                                               ,basefreq$c[5],basefreq$g[5])),collapse =""),
                           collapse ="",sep=""
      )
      
      write.fasta(DNA_sequence, 
                  line_header, 
                  file.out=paste(YOUR_Set,"AnalogousRNAs",iii,".fasta", sep=""), 
                  open = "a", 
                  nbchar = 60, 
                  as.string = TRUE)
    }}}

#Filtering of CDS for completeness

library(seqinr)
library(sfsmisc)
YOUR_CDS<-""  #include folder

CDS<-read.fasta(YOUR_CDS)
cleanSeqs<-c()
for (i in 1:length(CDS)){
  if(((CDS[[i]][[1]]==c("a")&
       (CDS[[i]][[2]]==c("t"))&
       (CDS[[i]][[3]]==c("g"))))&(
         (CDS[[i]][[((length(CDS[[i]]))-2)]]==c("t")&
          (CDS[[i]][[(length(CDS[[i]]))-1]]==c("a"))&
          (CDS[[i]][[(length(CDS[[i]]))]]==c("g")))|
         (CDS[[i]][[(length(CDS[[i]]))-2]]==c("t")&
          (CDS[[i]][[(length(CDS[[i]]))-1]]==c("a"))&
          (CDS[[i]][[(length(CDS[[i]]))]]==c("a")))|
         (CDS[[i]][[(length(CDS[[i]]))-2]]==c("t")&
          (CDS[[i]][[(length(CDS[[i]]))-1]]==c("g"))&
          (CDS[[i]][[(length(CDS[[i]]))]]==c("a")))
       )&(is.whole((length(CDS[[i]]))/3)==TRUE)){cleanSeqs<-c(cleanSeqs,i)}
}

CDS<-CDS[cleanSeqs]

write.fasta(sequences=CDS, 
            names=names(CDS), 
            file.out=paste(YOUR_CDS,".cleaned.fasta", sep=""), 
            open = "w", 
            nbchar = 100)


#Calculation of the P-CHS
#The results of the tapir target prediction output are written to a file and saved to a directory (YOUR_directory)
#Do not delete or move your sRNA sets

library(seqinr)
library(sfsmisc)
YOUR_directory<-""
YOUR_TAPIR_results<-"" #The file containing the TAPIR output
YOUR_protocol<-""
YOUR_filtered_CDS<-"" #paste(YOUR_CDS,".cleaned.fasta", sep="")


file.create(YOUR_protocol)

cleaned_CDS<-read.fasta(YOUR_filtered_CDS)


all.cds.uco<-uco(unlist(getSequence(cleaned_CDS)), frame = 0, index =  c("eff", "freq", "rscu"), as.data.frame = TRUE,
                 NA.rscu = NA)
write.csv(all.cds.uco,paste0(YOUR_directory,"cds.uco.csv"))


checkedi<-c()

vi<-YOUR_TAPIR_results

print(vi)
write(paste("target file",vi),file=YOUR_protocol,append=TRUE)
tapir_results1<-read.csv(paste0(YOUR_directory,YOUR_TAPIR_results),header=T, sep="\t")
tapir_results1$X..Search.options<-as.character(tapir_results1$X..Search.options)

miRNA_seqs<-read.fasta(gsub(".* ","",tapir_results1$X..Search.options[1]))

tapir_results11<-subset(tapir_results1, !grepl("^#",tapir_results1$X..Search.options))
tapir_results12<-subset(tapir_results11, !grepl("//",tapir_results11$X..Search.options))
tapir_results12$X..Search.options <- gsub(".*[ ]","",tapir_results12$X..Search.options)
tapir_results14<-data.frame(matrix(unlist(tapir_results12), ncol=14, byrow=T))
names (tapir_results14) <- c("miRNA","target","score","mfe_ratio","start","seed_gap","seed_mismatch"
                             ,"seed_gu","gap","mismatch","gu","miRNA_3'","aln","target_5'")

tapir_results14$Gene<-gsub("\\..*","",tapir_results14$target)


tapir_results14$miRNA<-as.character(tapir_results14$miRNA)
tapir_results14$target<-as.character(tapir_results14$target)
tapir_results14$score<-as.numeric(as.character(tapir_results14$score))
tapir_results14$mfe_ratio<-as.numeric(as.character(tapir_results14$mfe_ratio))
tapir_results14$start<-as.numeric(as.character(tapir_results14$start))
tapir_results14$seed_gap<-as.numeric(as.character(tapir_results14$seed_gap))
tapir_results14$seed_mismatch<-as.numeric(as.character(tapir_results14$seed_mismatch))
tapir_results14$seed_gu<-as.numeric(as.character(tapir_results14$seed_gu))
tapir_results14$gap<-as.numeric(as.character(tapir_results14$gap))
tapir_results14$mismatch<-as.numeric(as.character(tapir_results14$mismatch))
tapir_results14$gu<-as.numeric(as.character(tapir_results14$gu))
tapir_results14$`miRNA_3'`<-as.character(tapir_results14$`miRNA_3'`)
tapir_results14$aln<-as.character(tapir_results14$aln)
tapir_results14$`target_5'`<-as.character(tapir_results14$`target_5'`)
tapir_results14$Gene<-as.character(tapir_results14$Gene)

for(xii in 1:length(tapir_results14$miRNA)){
  if(tapir_results14[xii,]$score==0){
    perf.match<-tapir_results14[xii,]
    perf.match$`miRNA_3'`<-toupper(c2s(getSequence(miRNA_seqs[perf.match$miRNA])[[1]][length(getSequence(miRNA_seqs[perf.match$miRNA])[[1]]):1]))
    perf.match$`target_5'`<-gsub("T","U",toupper(c2s(comp(gsub("u","t",getSequence(miRNA_seqs[perf.match$miRNA])[[1]][length(getSequence(miRNA_seqs[perf.match$miRNA])[[1]]):1])))))
    tapir_results14[xii,]<-perf.match
  }
}

print(paste("all",length(tapir_results14$miRNA)))
write(paste("all",length(tapir_results14$miRNA)),file=YOUR_protocol,append=TRUE)

checkedi<-c(1:length(tapir_results14$miRNA))[tapir_results14$`target_5'`==""]
if(length(checkedi)>=1){
  print("No target sequence for interaction:")
  print(checkedi)
  stop()
}
dup14<-tapir_results14[,c(1,14,15)]
tapir_results14<-tapir_results14[!duplicated(dup14),]

print(paste("duplicated",length(tapir_results14$miRNA)))
write(paste("duplicated",length(tapir_results14$miRNA)),file=YOUR_protocol,append=TRUE)

cleaned_CDS_targets<-cleaned_CDS[getName(cleaned_CDS)%in%unique(tapir_results14$target)]

p.same.vec<-c()
p.bigger.vec<-c()
p.smaller.vec<-c()
print("2")
for (i in 1:length(tapir_results14[,1])){
  bulge.gene<-c()
  if(grepl("-",tapir_results14[i,14])){
    bulge.gene<-c(1:length(s2c(tapir_results14[i,14])))[grepl("-",s2c(tapir_results14[i,14]))]
  }
  
  
  #select codons overlapping whole sRNA target region
  sRNAstart<-tapir_results14[i,5]
  sRNAframestart=sRNAstart
  while (is.whole((sRNAframestart-1)/3)==F){sRNAframestart=sRNAframestart-1}
  sRNAend=sRNAstart+nchar((tapir_results14[i,13]))-1
  if(length(bulge.gene)>=1){sRNAend=sRNAend-length(bulge.gene)}
  sRNAframeend=sRNAend
  while (is.whole((sRNAframeend)/3)==F){sRNAframeend=sRNAframeend+1}
  if(sRNAframeend>
     getLength(cleaned_CDS_targets[tapir_results14[i,2]])){
    sRNAframeend=getLength(cleaned_CDS_targets[tapir_results14[i,2]])
  }
  if(sRNAframestart<=0){sRNAframestart=1}
  cds.target.seq<-  getFrag(cleaned_CDS_targets[as.character(tapir_results14[i,2])],
                            sRNAframestart,sRNAframeend,name=paste(i,"target.region"))
  if(!all(unlist(getSequence(cds.target.seq))%in%c("a","c","g","t"))){
    
    p.same.vec<-c(p.same.vec,NA)
    p.bigger.vec<-c(p.bigger.vec,NA)
    p.smaller.vec<-c(p.smaller.vec,NA)
    print(paste0("non-ACGT in CDS of interaction: ",i))
    next}
  
  p.same<-c()
  p.bigger<-c()
  p.smaller<-c()
  no.cod<-((length(getSequence(cds.target.seq[[1]])))/3)
  permutations<-  expand.grid(rep(list(0:3),no.cod))
  perm.data<-data.frame(p.0=rep(c(0),no.cod),
                        p.1=rep(c(0),no.cod),
                        p.2=rep(c(0),no.cod),
                        p.3=rep(c(0),no.cod),
                        sim.perm=rep(c(0),no.cod))
  for (ii in 0:((length(getSequence(cds.target.seq[[1]]))-3)/3)){
    cod.on<-c2s(getSequence(cds.target.seq)[[1]][(ii*3+1):((ii+1)*3)])
    tar.seq.cod<-all.cds.uco[which(all.cds.uco$codon==cod.on),]
    tar.seq.aa<-all.cds.uco[which(all.cds.uco$AA==tar.seq.cod$AA),]
    tar.seq.aa<-tar.seq.aa[order(-tar.seq.aa$RSCU),]
    tar.seq.aa$pos<-(1:length(tar.seq.aa$RSCU))
    sRNAframestart.cod<-sRNAframestart+ii*3
    sRNAseq<-s2c(gsub("u","t",(tolower((tapir_results14[i,12])))))
    if(length(bulge.gene)>=1){sRNAseq=sRNAseq[-bulge.gene]}
    sRNAseq.comp<-comp(sRNAseq)
    sRNAseq.comp[is.na(sRNAseq.comp)] <- "y"
    tar.sRNA.offset<-sRNAstart-sRNAframestart
    sRNA.seq.cod.pos<-((ii*3)+1-tar.sRNA.offset):(((ii+1)*3)-tar.sRNA.offset)
    sRNA.seq.cod<-c()
    for (iii in sRNA.seq.cod.pos){
      if(iii<=0){
        sRNA.seq.cod<-c(sRNA.seq.cod,"x")}
      else{
        sRNA.seq.cod<-c(sRNA.seq.cod,sRNAseq.comp[iii])
        sRNA.seq.cod[is.na(sRNA.seq.cod)]<-"x"
      }
    }
    similarity<-c()
    for(iv in 1:length(tar.seq.aa$codon)){
      compare<-c()
      for(v in 1:3){
        compa<-grepl(sRNA.seq.cod[v],s2c(as.character(tar.seq.aa$codon[iv]))[v])
        
        compare<-c(compare,compa)
      }
      simi<-sum(as.numeric(compare))
      similarity<-c(similarity,simi)
    }
    tar.seq.aa$similarity<-similarity
    tar.seq.aa$RRSCU<-tar.seq.aa$RSCU/length(tar.seq.aa$RSCU)
    tar.cod<-tar.seq.aa[which(tar.seq.aa$codon==cod.on),]
    perm.data$p.0[ii+1]<-sum(tar.seq.aa[which(tar.seq.aa$similarity==0),]$RRSCU)
    perm.data$p.1[ii+1]<-sum(tar.seq.aa[which(tar.seq.aa$similarity==1),]$RRSCU)
    perm.data$p.2[ii+1]<-sum(tar.seq.aa[which(tar.seq.aa$similarity==2),]$RRSCU)
    perm.data$p.3[ii+1]<-sum(tar.seq.aa[which(tar.seq.aa$similarity==3),]$RRSCU)
    perm.data$sim.perm[ii+1]<-tar.cod$similarity
    tar.main<-tar.seq.aa[1,]
    
  }
  permutations<-transform(permutations, sum=rowSums(permutations))
  permutations2<-data.frame(matrix(NA, nrow = length(permutations[,1]), ncol = no.cod))
  for(x in 1:(no.cod)){
    permutations2[,x] [permutations[,x]==0]<-perm.data$p.0[x]
    permutations2[,x] [permutations[,x]==1]<-perm.data$p.1[x]
    permutations2[,x] [permutations[,x]==2]<-perm.data$p.2[x]
    permutations2[,x] [permutations[,x]==3]<-perm.data$p.3[x]
  }
  permutations2$sum<-permutations$sum
  sum.sim<-sum(perm.data$sim.perm)
  x.p = apply(permutations2[,c(1:no.cod)], MARGIN = 1, FUN = prod)
  permutations2$p<-x.p
  p.same<-sum(permutations2[which(permutations2$sum==sum(perm.data$sim.perm)),]$p)
  p.bigger<-sum(permutations2[which(permutations2$sum>sum(perm.data$sim.perm)),]$p)
  p.smaller<-sum(permutations2[which(permutations2$sum<sum(perm.data$sim.perm)),]$p)
  
  if(!(round(p.same+p.bigger+p.smaller,digits=9)==1)){
    print("ERROR: total probability =/= 1")
    print(i)
  }
  
  p.same.vec<-c(p.same.vec,p.same)
  p.bigger.vec<-c(p.bigger.vec,p.bigger)
  p.smaller.vec<-c(p.smaller.vec,p.smaller)
  
}

tapir_results14$p.same<-p.same.vec
tapir_results14$p.bigger<-p.bigger.vec


results1<-tapir_results14
results1$PCHS<-results1$p.bigger+results1$p.same

print("PCHS calculated")

write.csv(results1,file=paste0(YOUR_directory,YOUR_TAPIR_results,"results_final.csv"))

#Localization of genomic regions targeted by sRNAs and extraction of specific FSTs

YOUR_genome<-""   #include folder
YOUR_VCFtools_FST_output<-"" #include folder. FSTs are stored by VCFtools as a file ending in ".weir.fst"
library(seqinr)
library(Biostrings)
library(biomaRt)

results1$end<-results1$start+(nchar(gsub("-","",results1$target_5.))-1)

#biomaRt needs to be set-up to support the specific organism
listMarts(host="https://plants.ensembl.org")
ensembl_Ath<-useEnsembl(biomart="plants_mart",
                        dataset="athaliana_eg_gene",
                        host="https://plants.ensembl.org")

mart_Ath=useMart(biomart="plants_mart",
                 dataset="athaliana_eg_gene",
                 host="https://plants.ensembl.org")


mart_Ath@biomart <- 'ENSEMBL_MART_ENSEMBL'

at_genome<-seqinr::read.fasta(YOUR_genome)

#seperation of genome into single chromosomes for process speed optimization

#if genome of interest is not A. thaliana chromosome identifiers need to be adjusted accordingly
chr1<-at_genome["NC_003070.9"] 
chr2<-at_genome["NC_003071.7"]
chr3<-at_genome["NC_003074.8"]
chr4<-at_genome["NC_003075.7"]
chr5<-at_genome["NC_003076.8"]
chrM<-at_genome["NC_001284.2"]
chrC<-at_genome["NC_000932.1"]

#seperate chromosomes are stored to a list
atChr<-list(chr1,
            chr2,
            chr3,
            chr4,
            chr5,
            chrM,
            chrC)

#data frames to store genomic positions
regions<-data.frame(CHROM=character(),
                    START=numeric(),
                    END=numeric())
regions1<-data.frame(CHROM=character(),
                     START=numeric(),
                     END=numeric())


#load your FSTs into a data frame
fst_set<-read.table(YOUR_VCFtools_FST_output,header=T,na.strings="-nan")

#seperation of FSTs into single chromosomes for process speed optimization
for(i in unique(fst_set$CHROM)){
  assign(paste0("fst_set_C",i),subset(fst_set,fst_set$CHROM==i))
}

#list to store FSTs. Each element will represent one line in your results
fst_list_set = list()

datalist = list()

z=1

#this loop will take some time to compute
for (i in 1:length(results1$miRNA)){
  #preparation of empty lists, vectors and values
  #biomaRt data
  bioM1<-data.frame(chromosome_name=character(), ensembl_transcript_id=character(),
                    transcript_start=character(),transcript_end=character(),cds_start=character(),
                    cds_end=character(), genomic_coding_start=character(),
                    genomic_coding_end=character(), strand=character())
  
  fst_vec_set<-c() #vector of FSTs
  start1<-c() #vector to store start of genomic regions
  end1<-c() #vector to store end of genomic regions
  nuc_vec<-c() #vector to store nucleotide sequence for internal check
  split=F #value to indicate if target region spans a junction
  
  #data acquisition
  start<-results1$start[i] #start of target region in transcript
  end<-results1$end[i] #end of target region in transcript
  trans<-results1$target[i] #transcript of interest
  
  #fetch genomic coordinates and transcript information from biomaRt
  bioM<-getBM(attributes=c("chromosome_name","ensembl_transcript_id","transcript_start","transcript_end",
                           "cds_start","cds_end",
                           "genomic_coding_start","genomic_coding_end",
                           "strand"),
              filters="ensembl_transcript_id",
              values=trans,
              mart=mart_Ath
  )
  chrom<-bioM[1,1] #chromosome
  chrom1<-chrom
  #mitochondrial and plastidal chromosomes are excluded from analysis, due to lack of data
  if(chrom=="Mt"){chrom1<-6
  next}
  if(chrom=="Pt"){chrom1<-7
  next}
  
  #relevant exons are selected and stored in "bioM1" 
  #and junction spanning target regions are identified
  for(ii in 1:length(bioM$chromosome_name)){
    if(is.na(bioM[ii,5])){next}
    if(start>=bioM$cds_start[ii]&end<=bioM$cds_end[ii]){
      bioM1<-bioM[ii,]
    }
    if(!(start>=bioM$cds_start[ii]&end<=bioM$cds_end[ii])&(
      (start>=bioM$cds_start[ii]&end>=bioM$cds_end[ii]&start<=bioM$cds_end[ii])|
      (end<=bioM$cds_end[ii]&start<=bioM$cds_start[ii]&end>=bioM$cds_start[ii]))){
      split=T
      bioM1<-rbind(bioM1,bioM[ii,])
    }
  }
  #regions for non-spanning transcripts are determined
  if(!split){
    #plus strand coordinates
    if(bioM1$strand==1){
      start1=start+bioM1$genomic_coding_start-bioM1$cds_start
      end1=end+bioM1$genomic_coding_start-bioM1$cds_start
      regions1<-data.frame(CHROM=chrom,
                           START=start1,
                           END=end1)
      #FSTs are stored in "fst_vec_set"
      fst_set_1<-get(paste0("fst_set_C",regions1$CHROM[1]))
      
      fst_set_2<-subset(fst_set_1,fst_set_1$POS>=regions1$START[1])
      fst_set_3<-subset(fst_set_2,fst_set_2$POS<=regions1$END[1])
      fst_vec_set<-c(fst_vec_set,fst_set_3$WEIR_AND_COCKERHAM_FST)
      #genomic regions are stored in "datalist"
      datalist[[z]] <- regions1
      z=z+1

    }
    #plus strand coordinates
    if(bioM1$strand==-1){
      start1=bioM1$genomic_coding_end-start+bioM1$cds_start
      end1=bioM1$genomic_coding_end-end+bioM1$cds_start
      regions1<-data.frame(CHROM=chrom,
                           START=end1,
                           END=start1)
      
      #FSTs are stored in "fst_vec_set"
      fst_set_1<-get(paste0("fst_set_C",regions1$CHROM[1]))
      
      fst_set_2<-subset(fst_set_1,fst_set_1$POS>=regions1$START[1])
      fst_set_3<-subset(fst_set_2,fst_set_2$POS<=regions1$END[1])
      fst_vec_set<-c(fst_vec_set,fst_set_3$WEIR_AND_COCKERHAM_FST)
      
      #genomic regions are stored in "datalist"
      datalist[[z]] <- regions1
      z=z+1

    }
    
    #genomic coordinates are compared to target sequence in the TAPIR results
    #plus strand
    seq1<-toupper(gsub(", ","",toString(unlist(seqinr::getSequence(atChr[[chrom1]]))[c(start1:end1)])))
    seq2<-gsub("U","T",gsub("-","",results1$target_5.[i]))
    #minus strand
    if(bioM1$strand==-1){
      seq1<-toupper(gsub(", ","",toString(unlist(seqinr::getSequence(atChr[[chrom1]]))[c(end1:start1)])))
      seq2<-gsub("U","T",gsub("-","",results1$target_5.[i]))
      seq1<-as.character(reverseComplement(DNAString(seq1)))}
    #if genomic sequence does not match the sequence of the TAPIR target sequence a error is reported
    #loop execution continues. Results are not valid!
    if(!seq1==seq2){print(paste0("problem: ",i))}
  }
  #regions for junction-spanning transcripts are determined
  if(split){
    bioM1<-bioM1[order(bioM1$cds_start),] 
    #plus strand coordinates
    if(bioM1$strand[1]==1){
      #selection of target region overlapping exons
      for(iii in 1:length(bioM1$chromosome_name)){
        if(bioM1$cds_start[iii]<=start&bioM1$cds_end[iii]<=end){
          start1=start+bioM1$genomic_coding_start[iii]-bioM1$cds_start[iii]
          end1=bioM1$genomic_coding_end[iii]
          #vector for internal error check is generated
          nuc_vec<-c(nuc_vec,(start+bioM1$genomic_coding_start[iii]-bioM1$cds_start[iii]):
                       (bioM1$genomic_coding_end[iii]))
        }
        if(bioM1$cds_start[iii]>=start&bioM1$cds_end[iii]>=end){
          start1=bioM1$genomic_coding_start[iii]
          end1=bioM1$genomic_coding_start[iii]-bioM1$cds_start[iii]+end
          #vector for internal error check is generated
          nuc_vec<-c(nuc_vec,(bioM1$genomic_coding_start[iii]):
                       (bioM1$genomic_coding_start[iii]-bioM1$cds_start[iii]+end)
          )
          
          
        }

        regions1<-data.frame(CHROM=chrom,
                             START=start1,
                             END=end1)
        #FSTs are stored in "fst_vec_set"
        fst_set_1<-get(paste0("fst_set_C",regions1$CHROM[1]))
        
        fst_set_2<-subset(fst_set_1,fst_set_1$POS>=regions1$START[1])
        fst_set_3<-subset(fst_set_2,fst_set_2$POS<=regions1$END[1])
        fst_vec_set<-c(fst_vec_set,fst_set_3$WEIR_AND_COCKERHAM_FST)
        
        datalist[[z]] <- regions1
        z=z+1
        
      }
    }
    #minus strand coordinates
    if(bioM1$strand[1]==-1){
      #selection of target region overlapping exons
      for(iii in length(bioM1$chromosome_name):1){
        if(bioM1$cds_start[iii]>=start&bioM1$cds_start[iii]<=end){
          #vector for internal error check is generated
          nuc_vec<-c(nuc_vec,(bioM1$genomic_coding_end[iii]+bioM1$cds_start[iii]-end):
                       (bioM1$genomic_coding_end[iii]))
          end1=(bioM1$genomic_coding_end[iii])
          start1=(bioM1$genomic_coding_end[iii]+bioM1$cds_start[iii]-end)
        }
        if(bioM1$cds_start[iii]<=start&bioM1$cds_end[iii]<=end){
          #vector for internal error check is generated
          nuc_vec<-c(nuc_vec,(bioM1$genomic_coding_start[iii]):
                       (bioM1$genomic_coding_start[iii]+bioM1$cds_end[iii]-start))
          start1=(bioM1$genomic_coding_start[iii])
          end1=(bioM1$genomic_coding_start[iii]+bioM1$cds_end[iii]-start)
        }
        regions1<-data.frame(CHROM=chrom,
                             START=start1,
                             END=end1)
        #FSTs are stored in "fst_vec_set"
        fst_set_1<-get(paste0("fst_set_C",regions1$CHROM[1]))
        
        fst_set_2<-subset(fst_set_1,fst_set_1$POS>=regions1$START[1])
        fst_set_3<-subset(fst_set_2,fst_set_2$POS<=regions1$END[1])
        fst_vec_set<-c(fst_vec_set,fst_set_3$WEIR_AND_COCKERHAM_FST)
        
        datalist[[z]] <- regions1
        z=z+1
        
      }
    }
    #genomic coordinates are compared to target sequence in the TAPIR results
    #plus strand
    seq1<-toupper(gsub(", ","",toString(unlist(seqinr::getSequence(atChr[[chrom1]]))[nuc_vec])))
    seq2<-gsub("U","T",gsub("-","",results1$target_5.[i]))
    #minus strand
    if(bioM1$strand[1]==-1){
      seq1<-as.character(reverseComplement(DNAString(seq1)))
    }
    #if genomic sequence does not match the sequence of the TAPIR target sequence a error is reported
    #loop execution continues. Results are not valid!
    if(!seq1==seq2){print(paste0("problem: ",i))}
    #junction spanning target regions and strand are reported
    #this is normal behaviour and no action is necessary
    print(paste0("split: ",i," strand: ",bioM1$strand[1]))   
  }
  #due to the long execution time of this loop progress is reported every 25 interactions
  #this is normal behaviour and no action is necessary
  if(i%%25==0){print(i)}
  
  #FSTs are stored in "fst_list_set". Each element in list corresponds to one row in TAPIR results.
  fst_list_set[[i]]<-fst_vec_set

}

regions2 = do.call(rbind, datalist)
#Test if all regions are stored in correct direction
#If FALSE something went wrong 
all((regions2$end-regions2$start)>=1)
#regions2 can be written to file to be used as regions file in BCF or VCFtools for downstream analyses
#file name needs to be specified
write.table(regions2,file="",quote=F,sep="\t",
            row.names=F,col.names=F)

#For ploting FSTs can be stored to a data frame with additional information 
results_fst<-data.frame(set=character(),
                         pchs=numeric(),
                         fst=numeric())

for(i in 1:length(fst_list_set)){
  results_fst_1<-data.frame(set=rep(results1$set[i],length(fst_list_set[[i]])),
                             pchs=rep(results1$PCHS[i],length(fst_list_set[[i]])),
                             fst=fst_list_set[[i]])

  results_fst<-rbind(results_fst,results_fst_1)
}

#"results_fst" still contains NAs and negative FSTs which needs accounting in downstream applications
#e.g.
fst_set_noNA<-fst_set[complete.cases(fst_set),]
fst_set_noNA_0<-fst_set_noNA
fst_set_noNA_0$WEIR_AND_COCKERHAM_FST[fst_set_noNA$WEIR_AND_COCKERHAM_FST<=0]<-0
